# Constructive Connectomics: how neuronal axons get from here to there using gene-expression maps derived from their family trees

**DOI:** 10.1101/2022.02.26.482112

**Authors:** Stan Kerstjens, Gabriela Michel, Rodney J. Douglas

## Abstract

During brain development, billions of axons must navigate over multiple spatial scales to reach specific neuronal targets, and so build the processing circuits that generate the intelligent behavior of animals. However, the limited information capacity of the zygotic genome puts a strong constraint on how, and which, axonal routes can be encoded. We propose and validate a mechanism of development that can provide an efficient encoding of this global wiring task. The key principle, confirmed through simulation, is that basic constraints on mitoses of neural stem cells—that mitotic daughters have similar gene expression to their parent and do not stray far from one another—induce a global hierarchical map of nested regions, each marked by the expression profile of its common progenitor population. Thus, a traversal of the lineal hierarchy generates a systematic sequence of expression profiles that traces a staged route, which growth cones can follow to their remote targets. We have analyzed gene expression data of developing and adult mouse brains published by the Allen Institute for Brain Science, and found them consistent with our simulations: gene expression indeed partitions the brain into a global spatial hierarchy of nested contiguous regions that is stable at least from embryonic day 11.5 to postnatal day 56. We use this experimental data to demonstrate that our axonal guidance algorithm is able to robustly extend arbors over long distances to specific targets, and that these connections result in a qualitatively plausible connectome. We conclude that, paradoxically, cell division may be the key to uniting the neurons of the brain.

**Author Summary:** The embryological development of each brain installs an essentially identical communication network between its cells that is roughly as complex as that between the billions of people living on Earth. Although vast scientific resources are currently applied to identifying the final pattern of connections, the *connectome*, there has until now been relatively little effort to answer the fundamental question of how this complex network across billions of neurons realized through the mitotic elaboration of the initial embryonic cell. The problem is sharpened by the constraints that construction of the network is limited by the information budget of the initial genome, and that it has no pre-existing address space for placing neurons and guiding axons. We explain how Biology can solve this problem by using the family tree of neurons to install a global space of molecular addresses, which axons can use to navigate from their source neuron to its relatives. We provide experimental evidence for this familial address space in gene expression patterns of the developing mouse brain, and demonstrate through simulation that the experimentally observed address space indeed supports global navigation to produce a qualitatively plausible default connectome.

## Introduction

A century of neuroanatomical studies attest that, while their detailed synaptic configurations may differ, the fundamental organization of neuronal types and their axonal projections are highly conserved within a species. Major resources of neuroscience are currently devoted to describing the species-specific patterns of connections, or *connectomes*, of brains [40, 52, 12, 49, 62, 35, 16, 44, 29]. These projects typically approach the connectome from a reductive point of view: Given a mature brain, extract the graph of its neural nodes and axonal edges. This paper approaches the connectome from an entirely different point of view: Given a few progenitor cells derived from the zygote, explain their elaboration into the stereotypically connected graph of the brain. We will call this approach *Constructive Connectomics*.

There are two aspects to this construction problem: The generation, from a few precursors, of the vast number and various types of neuron that comprise the brain; and the process whereby these neurons then connect to one another. The process of generation is relatively well understood [42, 36, 43]. Undifferentiated progenitors who have inherited their genome from the zygote, undergo successive rounds of mitosis resulting in a exponentially large cell mass. At each division the mother cell gives rise to two daughters whose gene expression, and consequently whose phenotype, may differ from their mother and from one another. Overall, the branched sequence of mitoses can be represented as a lineage tree with undifferentiated progenitors at its root, and fully differentiated neurons at its leaves. Although some cell types do actively migrate, the cells of the growing mass generally maintain their location relative to one another. However, differentiated neuronal cells do give rise to excrescences tipped by growth cones, and these cones migrate away from their cell while drawing out an axon in their wake, actively searching for and then connecting to remote target neurons. Typically, the growth cone(s) will branch many times during this search, so generating a highly arborized axon that maps the source neuron through brain space to its many targets.

In vertebrates the fundamental wiring of the brain is established before birth, and occurs in near informational isolation from the external world. Therefore, the information required for stereotypical axonal guidance must be contained in the brain’s precursor cells, and hence is limited by the roughly 1GB information capacity of the original zygote^∗^. This amount is many orders of magnitude too small to explicitly encode sequences of receptor configurations for billions of neurons [64, 61, 26]. A naive encoding of this source-to-target connection matrix would require at least 10TB for a mouse brain, excluding the additional information required for detailed and staged axonal routing. The compression of the connectome into a 1GB genome implies that neuronal progenitors encode axon trajectories through brain space more efficiently than the naive approach. This raises the important question addressed here, of how neuronal progenitors encode and express the complex connections of the brain, and how this process is orchestrated through development (Figure 1).

**Figure 1:**
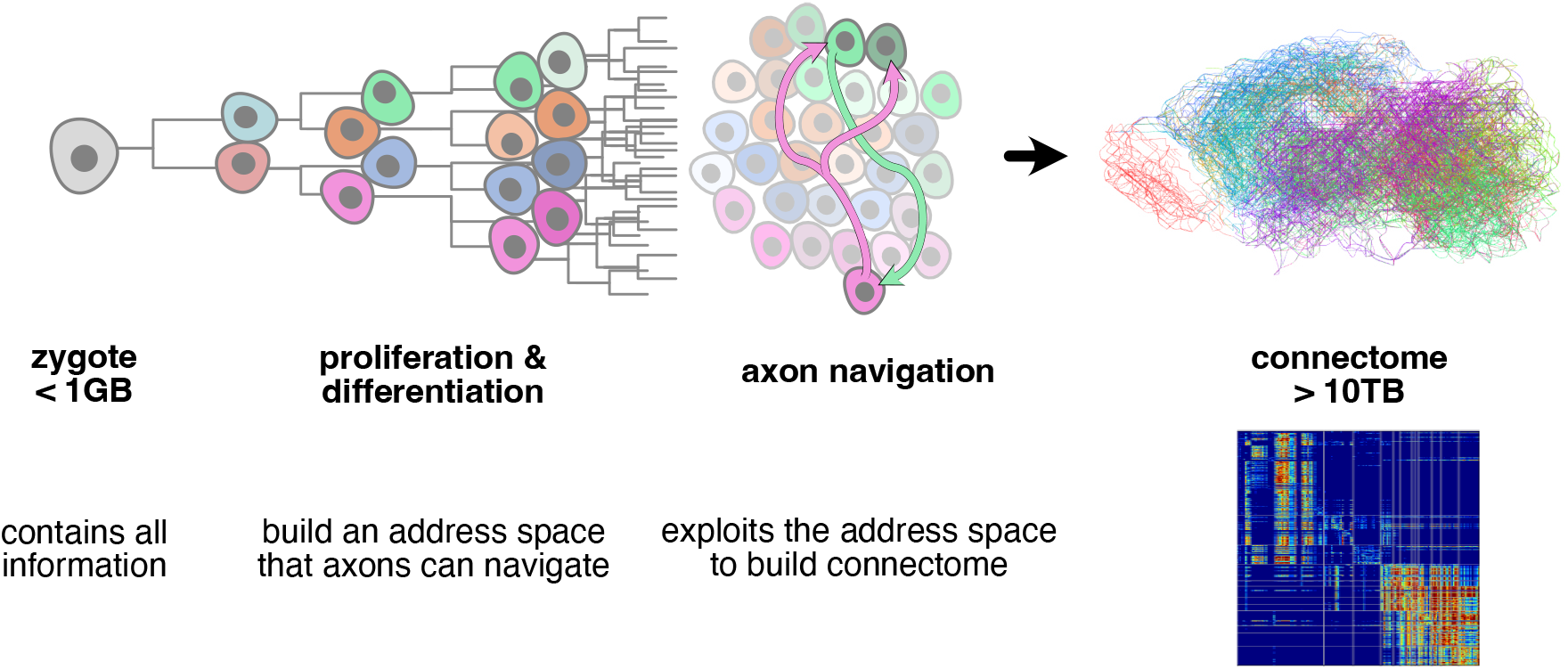
The connectome is the result of a constructive process that starts ultimately with the zygote, and involves the two aspects of first generating a mass of cells with various types, and then routing axons through this mass to their proper targets. An observer’s description of the resulting detailed mouse connection matrix (right bottom) takes at least 10TB to encode. However, as development occurs largely in isolation, all instructions to construct this connectome must fit into the 1GB of genetic material of the zygote. This implies that neural progenitors have efficient methods for expanding the highly compressed wiring instructions into axonal trajectories. To do this, they need to, as they proliferate and differentiate, install a space of molecular addresses that axons can exploit for navigation.

Growth cone guidance is crucial to the developing brain’s construction of its complex neuronal information processing circuits [6, 5, 50]. This claim is far from new: In his original description of axonal extension over a century ago, Cajal recognized that the cones are the agents of the neuronal circuit organization, and suggested that they are attracted to their targets by chemotaxis [7]. Later, Sperry elaborated that idea in his Chemoaffinity Hypothesis, whereby axons have differential markers; target cells have matching markers; markers are the result of cellular differentiation; and axons are actively directed by these markers to establish their specific connections [31, 48]. These tenets are now widely accepted, and there is by now broad experimental evidence that growth cones navigate by following gradients of molecular cues in their surrounding tissue [17, 5]. At decision points along their route, growth cones change their receptors to tune into a different molecular signal, and so change direction [32]. Long-range projections are achieved by growth cones passing through long sequences of growth cone receptor configurations [50].

Although these local molecular guidance mechanisms are relatively well understood, the global questions of just how these sequences and cues are encoded and deployed in both the navigating axons and navigated tissue (i.e., Sperry’s claim that they are installed by cellular differentiation) have been largely neglected. Our work addresses these issues and offers a principled account of how the connectome can be constructed within the zygote’s information budget.

Our working hypothesis is that the mitotic lineage tree induces a global hierarchical map over the developing brain. This map consists of nested regions of brain, each consisting of the progeny of a small definite population of progenitors whose profile of expression over multiple genes is maintained in the average expression profile of their collected progeny.

We propose that by successive differentiative mitoses of development, cells implicitly obtain unique gene expression addresses that encode their respective mitotic lineages (Figure 2). Then, by inverting the developmental program of their differentiation to revisit ancestral expression states, axons could effectively traverse the global family lineage tree by re-generating specific sequences of growth cone receptor configurations—each configuration seeking a region’s ancestral expression profile—that lead to long-range targets, which are in effect their mitotic cousins. That is, the sequence of intermediate targets between two neurons is implicit in their relative expression addresses, and requires no additional genetic encoding.

**Figure 2:**
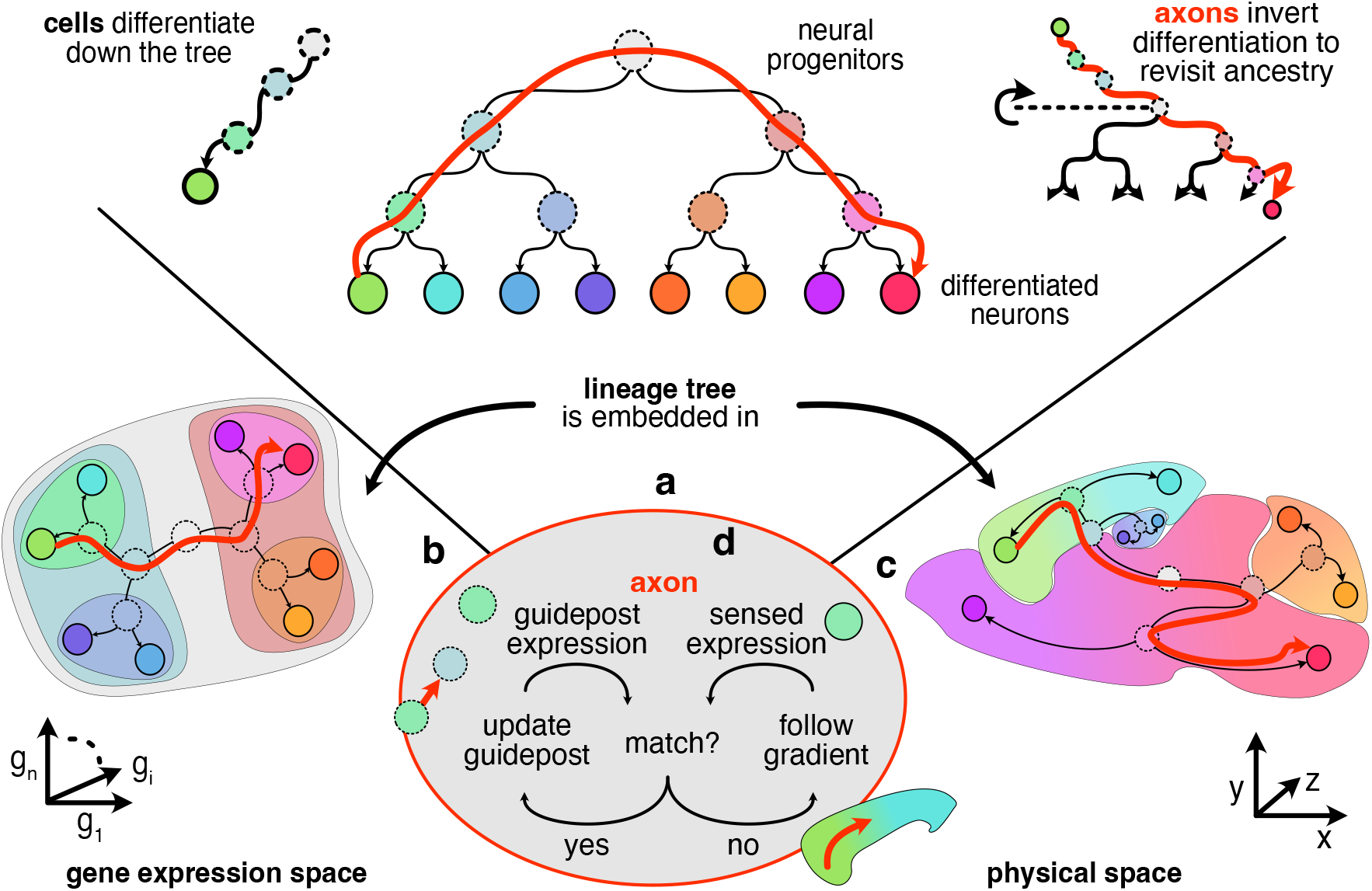
As progenitors divide they progressively differentiate their gene expression until they reach their post-mitotic neuronal states at the leaves of the lineage tree (**a**: top left). Constraints on mitosis (see text) embed the global neuronal lineage tree (**a**) into both gene expression (**b**) and physical space (**c**), so that cells of related differentiation have similar expression profiles (similar colors) and are nearby one another physically. Consequently, a trajectory from one leaf node to another through the lineage tree (**a**: red arrow) often corresponds to an unbroken trajectory through both gene expression space (**b**: red arrow) and physical brain space (**c**: red arrow). An axon navigates by inverting its source neuron’s instance of the global genetic differentiation program (**a**: top right). This inversion generates a sequence of expression profiles that correspond to ancestral states and so act as guidepost profiles. **d** The axonal branch configures its growth cone to match the sensed expression to the internally generated expression, and so moves to the direction that improves that match. When the match can no longer be improved by moving, the axon updates the its internal state to the next ancestor, and repeats. If the match between internal and external expression can be improved by moving into multiple different directions, or by transitioning to multiple different states, the single axonal branch is split into two new branches that continue to execute the same algorithm, but whose independent states may subsequently diverge. When an axonal branch arrives at a leaf state, both in expression and physical space, navigation of that branch is complete and local synapses are formed.

This guidance scheme requires that the lineage tree have a dual embedding in both expression and physical space (Figure 2). Through simulation, we show that this requirement is satisfied under rather simple conditions: That the mitotic daughters have gene expression similar to their parent; and, that the daughters do not usually migrate far from one another.

We find clear experimental evidence for our hypothesis in the *in situ* gene expression atlas published by the Allen Institute for Brain Science (ABI) [51], which provides voxelated spatial expression data of *∼*2000 developmentally relevant genes throughout the brain, extending from embryonic (E) day 11.5 to postnatal (P) day 56. Our analysis of their data reveals that the expression covariance of sets of randomly selected genes pattern the developing mouse brain on multiple spatial scales. These hierarchical patterns of expression involve the entire brain and spinal cord, transcend neuroanatomical boundaries, and are consistent over the available data. Furthermore, detailed simulations of our proposed guidance process on the ABI gene expression data confirm that axons can use it to robustly navigate over long distances to specific targets, as shown schematically in Figure 2.

We begin by describing in detail our overall concept, which provides the rationale for our analyses of experimental data, and for our simulations, presented in Results and Discussion. We conclude that the fundamental wiring of brain can be compactly encoded and expressed through the mitotic lineage implied by the genetic code of its embryonic stem cells. Thus, the connectome and its functioning can be more readily understood in terms of the global mechanisms that generate it (constructive connectomics), rather than from interpretation of the final wiring diagram (reductive connectomics), just as inspecting source code is more revealing of principles of operation than inspecting the compiled program.

## Rationale

The brain is an organized aggregation of billions of cells that are generated by many successive mitotic divisions of relatively few stem cells. Each of these stem cells is the root of a mitotic lineage tree that describes the branched sequence of mitoses beginning in that root and terminating in a population of post-mitotic leaf cells. Tracing these lineage trees experimentally, and understanding the relationship between the underlying cell states and their transitions during mitosis is a field of active research [23, 54, 53]. However, instead of pursuing the biological detail of these lineages, we explore an overarching statistical question: Could the sequence of mitoses impose an implicit order on the expression patterns of the leaf cells? And conversely, can the statistics of gene expression across a population of leaf cells offer an estimate of their shared lineage tree? The following section and Figure 3 offers an argument that both these properties are true, and lay a foundation for interpreting the experimental results and explanatory simulations that will follow in Results.

**Figure 3:**
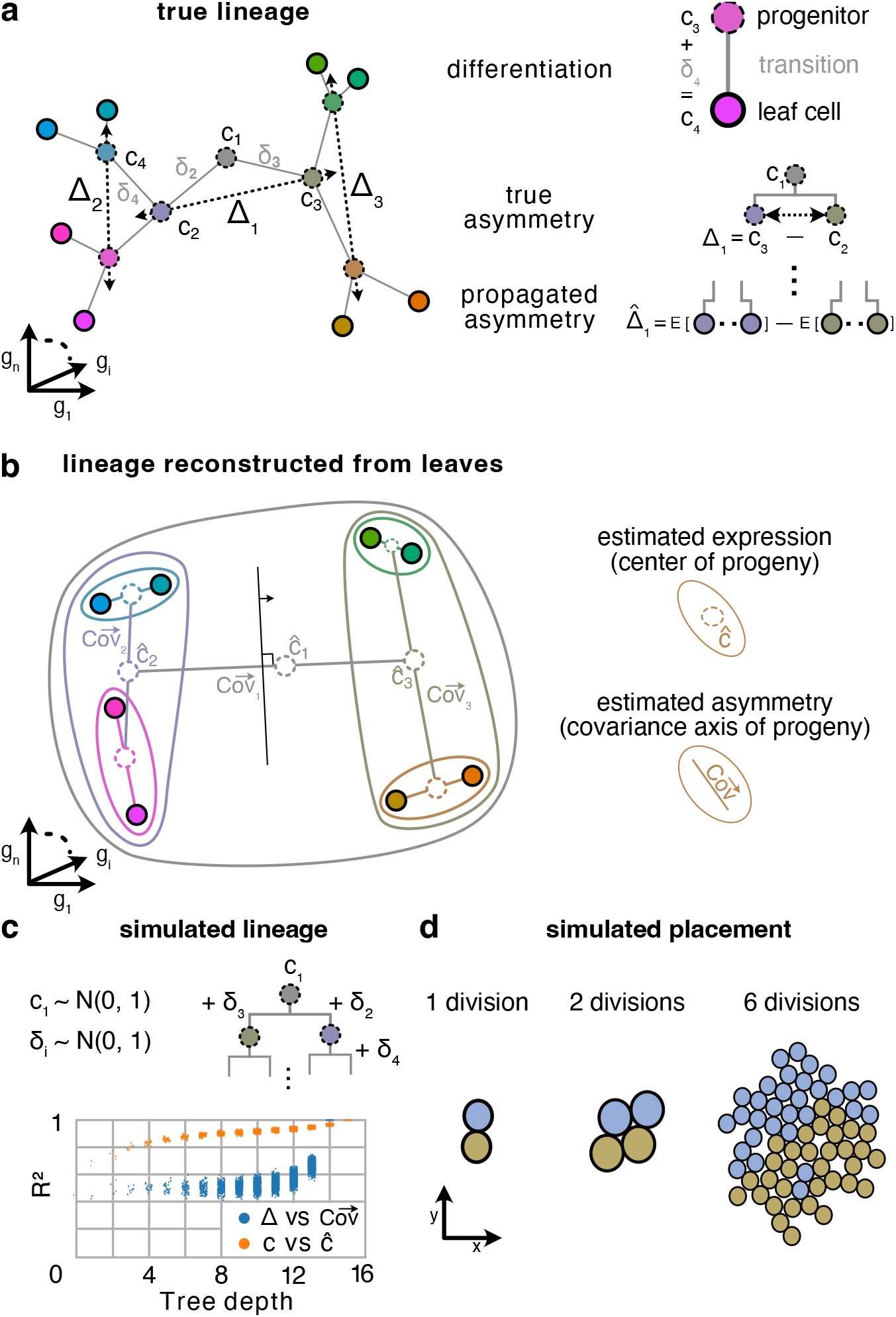
**a** Cells are points positioned in high-dimensional expression space, where each axis represents the expression of one gene. Here, this high-dimensional space is reduced to 2D dimensions for plotting purposes, so that their 2D distance approximates their high-dimensional distance. In our division model, the differential expression between a parent cell *c*_1_ and its daughters *c*_2_, *c*_3_ is a normally distributed random vector representing the genetic state transition from parent to daughter, denoted *δ*_2_ = *c*_2_ - *c*_1_. (Here we use the division of the root progenitor 1 as a running example for any division.) The differential expression between two siblings, which we call the parent’s asymmetry, is denoted .Δ_1_ = *c*_3_ - *c*_2_ = *δ*_3_ - *δ*_2_. As a result, the correlation in gene expression between two cells reflects their distance through the lineage tree. (See **c** for verification of this process by numerical simulation.) **b** The expression of a progenitor can be estimated as the mean expression over its leaf progeny; and the asymmetry of a progenitor can be measured as the main axis of variance across its progeny. The diagram shows only the leaves of the lineage tree show in **a**—they have identical positions in embedded expression space. Each nested contour encloses the progeny of a progenitor; lines within the countour indicate the main axis of variance across the enclosed progeny; and dotted circles the average expression across the progeny. The sets of progenies for individual progenitors can be obtained by iteratively splitting the progeny along their main axis of variance, so with a decision boundary (black line with arrow) orthogonal to this axis. **c** Numeric simulation of expression profiles induced by our division model, and subsequent reconstruction of expression profiles and mitotic asymmetries from the leaves of the simulated tree. The root expression *c*_1_ is drawn from a normal distribution with zero mean and unit variance. The expression profiles of other cells are generated recursively by adding differential expression patterns *δ*_*i*_, which are also normally distributed. (All random number are drawn independently.) The determination (squared correlation) was measured between between the true and reconstructed asymmetries (blue), and true and reconstructed expressions (orange). **d** Progenies group naturally in brain space according to their ancestry. Shown is a 2D simulation of growing tissue, started from a single root, only constrained to not detach from one another and not pass through each other.

### Familial Address Space Model

First we consider how and why the statistics of differential gene expression between mitotic siblings should be detectable across the brain as a map-like spatial hierarchy of gene expression covariance (Figure 3).

Our model for the gene expression around cell division is as follows. The life of each cell *i* starts as it is born from its mitotic mother, ends when it divides into its two mitotic daughters, and has an expression profile *c*_*i*_ over all genes and averaged over its lifetime. The lineage tree is rooted in progenitor cell 1, whose expression profile is *c*_1_. On mitosis, this parent cell splits into two daughter cells, 2 and 3, whose expression profiles become *c*_2_ = *c*_1_ + δ_2_, and *c*_3_ = *c*_1_ + δ_3_ respectively. Just so, the expression profile of every cell *i* in the lineage tree can be understood as the expression profile of its parent, plus a (positively or negatively signed) differential profile *δ*_*i*_ that summarizes the complex dynamics of gene regulation. The injection of these changes through mitosis is recursive, so that the gene expression of every cell is the accumulation of all its ancestral profiles, e.g., *c*_4_ = *c*_2_ + *δ*_4_ = (*c*_1_ + *δ*_2_)+ *δ*_4_.

Consider now the implications of these *δ*’s for the measurement of covariance in gene expression across populations of cells. The mitosis of progenitor 1 induces an expression asymmetry Δ_1_ = *c*_2_ - *c*_3_ = *δ*_2_ - *δ*_3_ between its daughters. This asymmetry propagates down the daughter’s two branches of the lineage, and is preserved across the two progenies as a quantity denoted as. 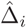 In other words, the parental mitosis injects a characteristic asymmetry that persists through development into the leaf cells (Figure 3**a**), and this signature could be detectable by an observer (Figure 3**b**). Every mitosis introduces such an asymmetry, so that the overall population of leaf cells, collectively, carry statistical evidence of the overall shape of the lineage. How can an observer extract this information? We argue (and demonstrate by simulations below) that lineage can be recovered by hierarchical decomposition of gene expression covariance.

Figure 3**b** sketches the method. Consider first the asymmetry Δ_1_ induced by *c*_1_. This asymmetry propagates down the lineage tree resulting in a trace. 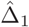 across all the leaf cells. We may estimate. 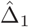 by measuring the direction of greatest variance of gene expression, denoted 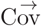 over the leaf offspring of 1. And, in general we expect that 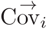, measured over the leaf cells of *i*, will be correlated with the original asymmetry Δ_*i*_ induced by mitosis of *c*_*i*_. The decomposition begins by measuring 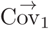 (across green cells). Then split these cells along 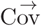 into two the daughter populations (red and blue); repeat 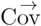 for each of these; and so on recursively. This hierarchical decomposition provides an estimate of the lineage tree.

The precise changes in gene expression *5* are, in general, unknown. But fortunately, detailed knowledge about the *δ*’s is not relevant for present purposes, only their general statistics. Figure 3**c** shows that. 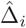 and .Δ_*i*_ are strongly correlated when (but not necessarily only when) the *δ*’s are independently and normally distributed.

Our Address Space Model asserts two principles. The first is that the profile of gene expression over multiple genes between a parent and daughter does not change on average. Although mitotic division may induce different gene expression profiles in the two daughters, both up-and down-regulation of any single gene are *a priori* equally likely. Thus, the expression profile over all genes averaged over both mitotic daughters will resemble the expression profile of the parent. This property (illustrated in Figure 3 **a**,**b**) is maintained recursively over successive cell divisions, such that the average expression over a given ancestor’s progeny resembles its own expression. It is this property that permits hierarchical decomposition: If we are able to select the progeny of a single progenitor while excluding cells from other branches of the lineage tree, then we could measure the asymmetry. 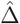 induced by the division of that progenitor by measuring 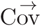 over the leaf progeny (see Figure 3**b**).

The second principle is that, also on average, the daughters of a mitosis do not stray too far from one another in 3D physical space. In this case, we expect that the mitotic lineage will be systematically organized across brain space, as seen in the clustering of each daughter’s progeny in Figure 3**d**. Therefore, mitotic asymmetries in gene expression (Figure 3**a**) are similarly organized in brain space (Figure 3**c**) and so encode a potential lineage address space. This associates a contiguous region in brain space with an ancestor in the lineage tree (larger regions correspond to earlier ancestors), and a trajectory through the lineage tree with a trajectory through brain space.

### Familial Guidance Model

First we describe our generic model (Figure 4) for the extension of a single axon arising from a source neuron in a cellular mass. Then, in Results we will report the application of this generic model to the voxelated case of the ABI data.

**Figure 4:**
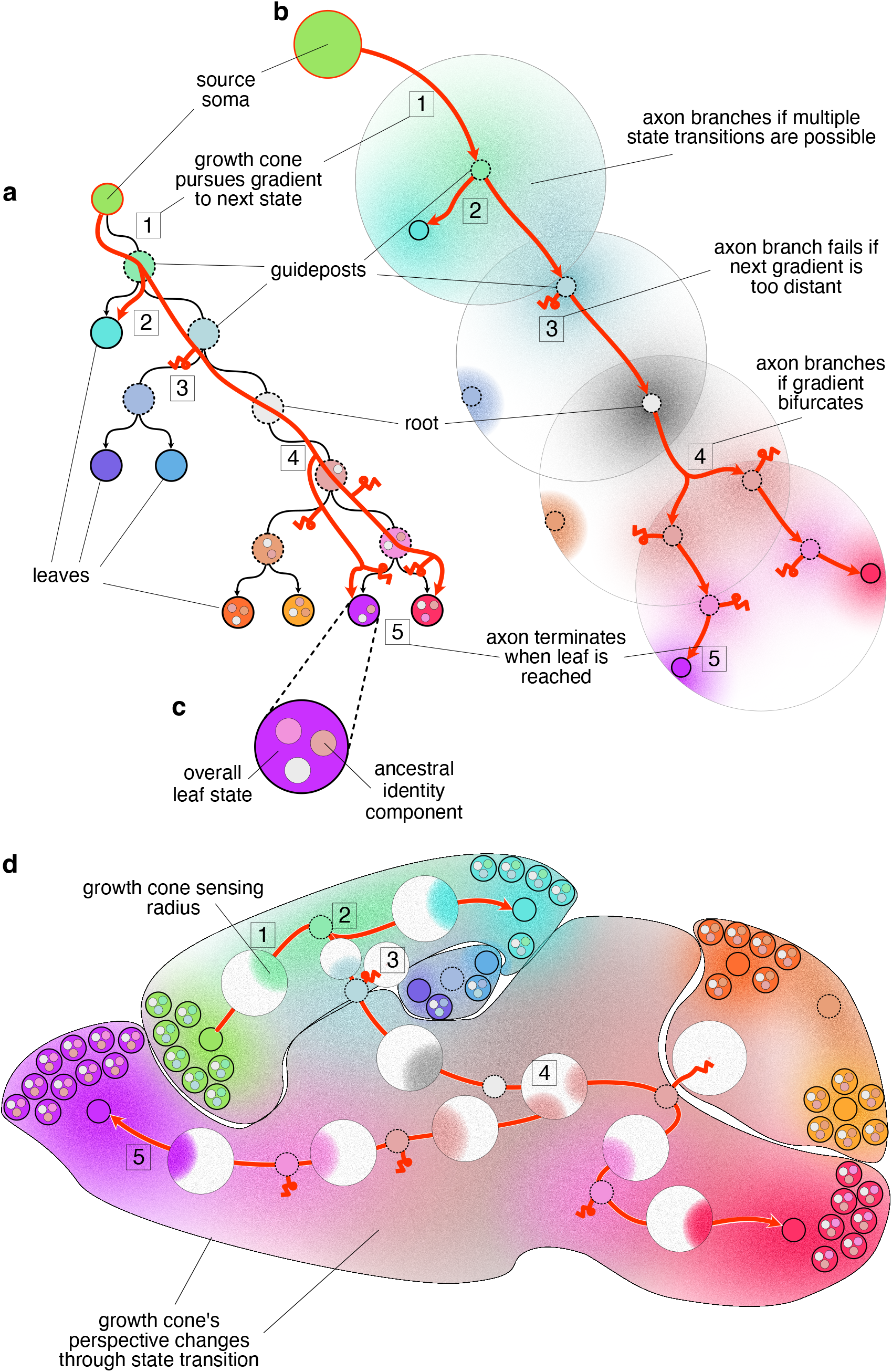
Navigation of an axon (red branching arrow) through the familial address space. Throughout the figure, similarity in color denotes similarity in gene expression profile. **a** The axon traverses the brain by traversing a sequence of familial states of the lineage tree that is implicit in its genome. The growth cone uses the sequence of familial states as successive search templates in brain space, and so navigates from a source leaf node to a number of target leaves. Familial states (colored circles) correspond to nodes of the encoded lineage tree. For purpose of explanation, the tree is hung from the leaf state corresponding to the axon’s source neuron, rather than from its root node as in 3b. Terminal states of leaf (existant) nodes have a solid circumference, while ancestral states in the interior of the tree have a dotted circumference. Transitions between states occur downward, along the arrowed arcs, beginning at the source leaf (red encircled) and ending at (some) other leaves. The original tree root can be recognized as the only state having two edges, rather than three (since the root progenitor has no mitotic parent). **b** Various decision scenarios that the axon encounters during traversal. Each familial state is characterized by a profile of gene expression, whose distribution across all cells peaks at one or more locations in brain space. The gradient of a state in the familial address space is the frequency of encountered cells that test positively for a familial state. By selecting a particular familial template, the growth cone tunes into the corresponding expression gradient and filters out the others. If the tuned gradient is in range, the growth cone follows it to arrive at one of that gradient’s peaks (case indicated by [1]). If the tuned gradient is not in range [3], the axonal branch of that growth cones fails. When the axon arrives at a peak, its growth cone tunes to the next downstream familial state, and so on, until a leaf state is found. If multiple downstream states are in range, the axon branches [2], with each branch tuned to one of the possible downstream states. The axon also branches if the gradient is bifurcated by a valley, so that the axon can follow an upward gradient in multiple directions [4]. Each branch pursues a different direction, but in this case they are tipped with growth cones in the same state (unlike the branches in scenario [2].) When a growth cone reaches a leaf state, guidance terminates [5]. **c** Cells have composite genetic identities, with one component (small inner circle) inherited from each ancestor state. The overall state of a leaf cell is the aggregation of these components (3). A growth cone can test whether a cell possesses a component by selecting the familial state template corresponding to that component, and then matching the internally produced gene expression to that of the tested cell. **d** Various regions of the brain correspond to branches of the mitotic lineage tree. Consequently, the regions are nested and each marked by the component of the genetic identity code corresponding to the common progenitor of the region.

The progeny of a given ancestor contain (on average) that ancestor’s gene expression signature. More-over, the expression profiles of cells are related (Figure 4, color gradients) by their ancestral sub-tree structure. Therefore, a particular axonal route through space from one cell to another is determined by the growth cone’s search over the successive expression signatures of the route through its ancestral lineage tree, which connects those two leaf cells (e.g., red path in Figure 4). Since the ancestral signatures are projected onto the leaf cells, the growth cone can navigate towards its target by maximizing the incidence of the successive signatures.

An axonal growth cone is instantiated by its source neuron. This growth cone extends its axon by moving in a direction that increases the match of its locally sensed expression with respect to search template. The cone selects as a search template the expression state of a node of the lineage tree. As explained above, these nodes represent ancestral expression patterns whose signatures can still be found in the current generation of leaf cells. Therefore the selection of a node as a template implies a search amongst local cells for that familial signature. The gradient of signature state in the familial address space is the frequency of encountered cells that test positively for that signature.

The lineage tree is implicitly encoded in every cell’s gene regulatory network, and the growth accesses templates by manipulating that network. Axonal growth and arborization results from successive optimization through growth cone movement, and replacement of the search template through genetic regulation in the axon, according to the following simple rules. The growth cone takes as its initial search template the leaf state of its source neuron. At each subsequent step of the search process the growth cone senses its external environment, and moves in a direction that satisfies its internal search template better than its current position. If there are other distinct directions that also increase satisfaction, then the growth cone divides and different axonal branches pursue each of those directions.

All branches of the same neuron are constrained by self-avoidance. In this sense, the paths in brainspace of a growing branch are irreversible, and their paths subject to race conditions. Additionally, before each step, the cone may replace its current search template with the expression state of any adjacent node in the lineage tree, so that the search will now be for a different, but closely related, familial template. However, such steps along the lineage tree are irreversible; and so this growth cone and its downstream branches will explore only those paths in brain-space and expression space that are coded in sub-tree sequences of expression templates. Growth cones terminate (or become dormant) when they can neither improve their template match, nor replace their template. The growth cone does distinguish between no gradient due to failure, and no gradient due to successfully reaching a peak. The cone will transition to the next possible state in both cases. If the cone continues to make progress with its search templates, it will ultimately reach a leaf state. If however, the local signals do not offer suitable gradients the growth cone will fail after exhausting its options.

In executing this search algorithm, the initial cone and its clones extend axons along all the routes in brain-space that offer contiguity in brain-space of familial expression patterns encoded in the lineage tree, and as a result create a particular repeatable axonal arborization from source to target leaf nodes.

## Results

We sought evidence of expression address maps by performing our hierarchical decomposition of covariances on experimentally measured gene expression data. We used the gene expression data published by the ABI in their Developing Mouse Brain Atlas [51]. Their data are provided as 3D grids of isotropic voxels. The expression energies of the *∼*2000 genes were measured by *in-situ* hybridization and take any non-negative value (see Methods).

Our prediction, leaning on the principles outlined above, was that the measured covariances should reveal a dual set of systematic patterns: a hierarchical ordering in expression space, and a nesting of region in brain space. Here, ‘expression space’ denotes the abstract multidimensional space in which the gene expression profile of a cell can be represented by a point; and ‘brain space’ or ‘space’ denotes physical 3D space. We will use ‘profile’ for a fixed relative expression of all genes within a cell or voxel, ‘pattern’ to denote an evident spatial regularity in gene expression, and ‘organization’ to denote the systematic order underlying these patterns.

### Global spatial expression hierarchy

We performed our hierarchical decomposition (described in the Concept and Methods sections) on the experimentally measured spatial gene expression, and consider to what extent that hierarchy could constitute an estimate of the mitotic lineage tree. Figure 5 shows the results for P28. Other time points from E11.5–P56 are shown in Figures S13–S20.

**Figure 5:**
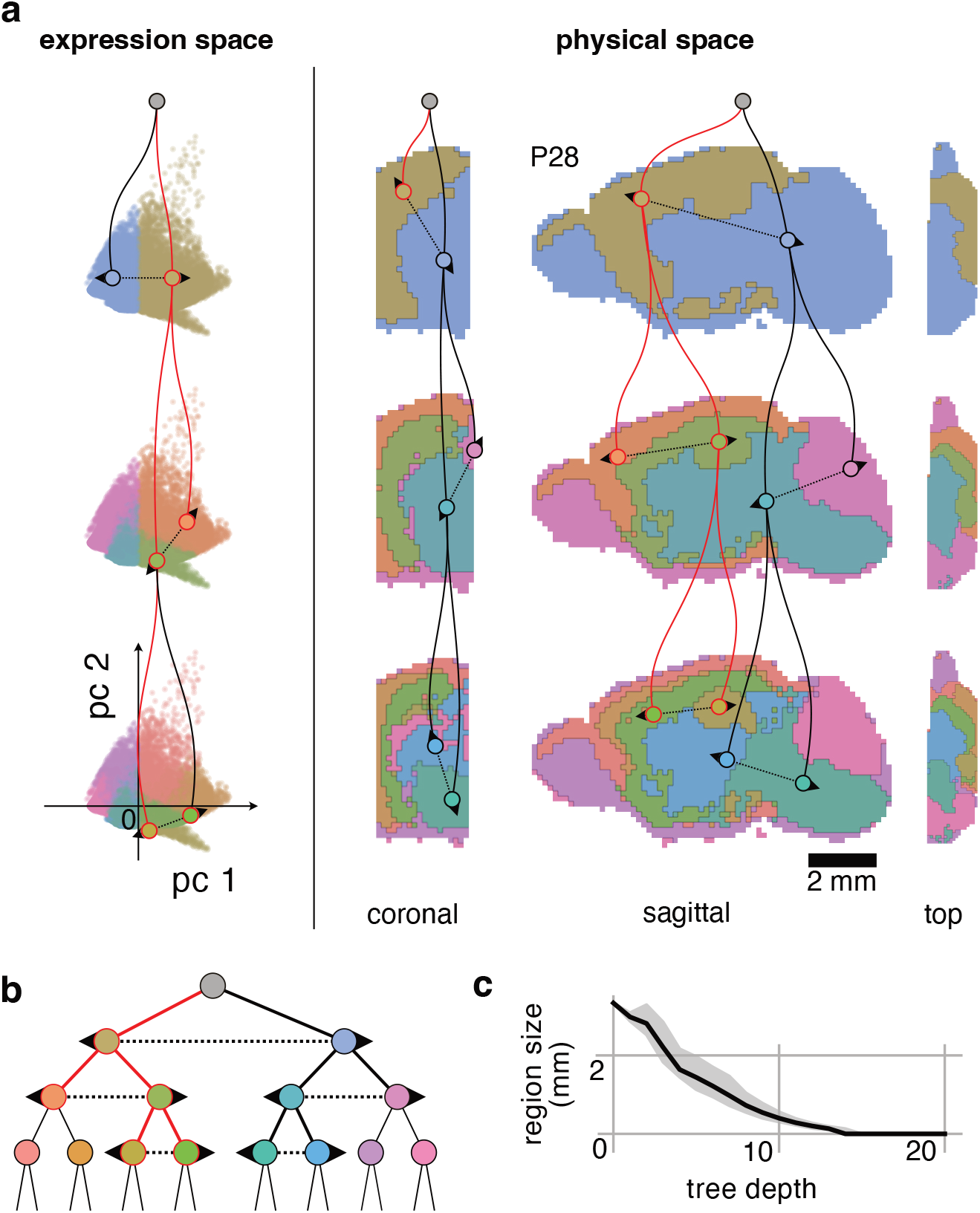
Hierarchical decomposition of covariance in gene expression space brain is mirrored by a matching decomposition in brain space. Here the results are for postnatal (P) day 28. The results from further time points can be found in the Supplementary figures. **(a) Expression** Hierarchical decomposition on the collection of voxels in expression space, independent of source location in brain. Decomposition is performed by measuring the first principal component of covariance; then sorting all voxels into two bins, (competing bins indicated by dotted arrows) based on their individual projection coefficients. This process is repeated recursively on each of the resulting bins, until a bin contains only a single voxel remains (Figure shows only the first 3 generations of the resulting hierarchy). **Physical Space** Same voxel bins and coloring, but voxels now positioned at their source locations in brain. Coronal and horizontal sections are shown: the color of each pixel indicates the most common bin in the occluded direction for that pixel. Horizontal section (labeled top) is drawn at a smaller scale. Multi-scale spatially coherent covariance patterns are present. Two example branches of the hierarchy are indicated with red and black curves. **(b)** Hierarchy of bins of the hierarchical decomposition. The bins are colored to represent the hierarchy: the parent bin has the average hue of the child bins. This coloring is applied throughout the paper. **(c)** Although regions are nested by construction (hierarchical decomposition), we quantified the extent to which the regions are also continuous by measuring their spatial spread (average distance from the region centroid) as a function of their depth in the hierarchy. At the root of the hierarchy the spatial spread covers the entire brain, and we expect that as the depth increases the spatial spread (i.e. the mean distance from the region centroid to the constituent voxels) decreases. To make the different time points and simulation comparable we present the spreads as a fraction of the root spread. The solid line indicates the median spread over all regions at that depth, and the gray area the first (below) and third (above) quartiles. As expected, the mean distance from the centroid decreases as the regions become more resolved at with depth.

The hierarchical decomposition is based exclusively on the gene expression covariance measured across unordered sets of voxels: The physical locations of voxels within the brain were not taken into account. However, when the voxels are now considered in their correct physical locations, we observe that the sets selected during the unordered hierarchical decomposition form nested and spatially continuous regions spanning brain space. Thus, the hierarchy of nested voxel sets found by our analysis in expression space parallels a hierarchy of spatially continuous regions in brain space, which is not entailed by the analysis.

To confirm that these patterns are due to intrinsic spatial organization of gene expression rather than being induced by the analysis, we performed the same decomposition on voxels with randomly shuffled gene expression values (see Methods). After shuffling, all spatial structure vanishes (Figures S13–S20).

We quantified the degree to which the hierarchy of asymmetries derived from gene expression projects into physical brain space by measuring the spatial spread of the voxel set. If the nested voxel sets form continuous regions in space beyond the first few generations, then the spread of their constituents should decrease over successive generations as the sub-populations become more resolved (and therefore smaller). Otherwise, if the constituent voxels are scattered across space, their spread would remain roughly constant. We find that indeed the spread consistently reduces with generation (Figure 5**c**).

In addition to spanning brain space, the patterns are consistent over time from E11.5 to P56. The temporal consistency cannot be measured directly, because there is no clear map between individual voxels of subsequent time points. To side-step this problem we instead project the voxels of one time point onto the hierarchy measured at another time point. In Figure 6 the voxels of all available time points are projected onto the hierarchy measured at P28.

**Figure 6:**
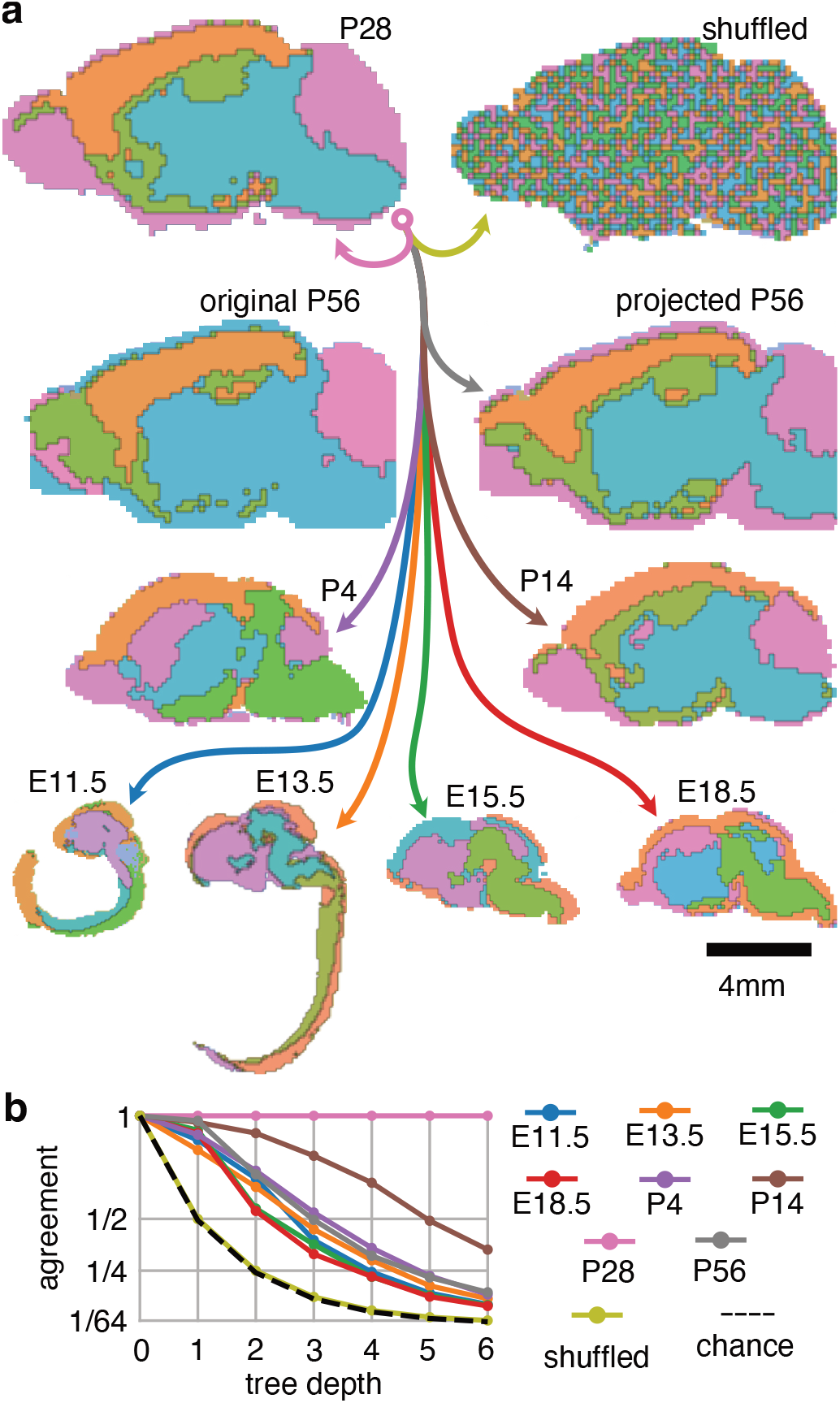
The root asymmetry measured at P28 is projected to the other available embryonic and postnatal developmental time points, and compared to the root asymmetry measured at the respective time point. **(a)** First division of the hierarchy, but the direction of variance used to sort the bins is derived from P28, rather than from the data of the time point itself (except Original P56). This temporally projected pattern only has small differences with the patterns derived from the original data (compare Original P56 to Projected P56). When the expression data is shuffled over voxels and genes, maintaining pooled expression statistics but destroying covariance structure, all spatial patterning disappears. Images are proportional to their actual brain sizes. **(b)** Quantification of the agreement between the original and projected hierarchy, measured as the proportion of voxels in matching bins, at different levels in the hierarchy. (Although the images in **a** are 2D, quantification is done on the 3D voxels.) The number of possible bins grows exponentially with tree depth, and so chance level decreases inverse-proportionally (dashed line), quantitatively verified by the shuffled case (yellow line). P28 projects onto itself, and is hence in perfect agreement. The other time points show an agreement consistently above chance. Consider that a mismatch at a shallow depth cannot be corrected at a deeper depth, and so mismatch can only accumulate.

To quantify the temporal consistency of the spatial patterns we measure how many voxels are sorted into the same hierarchical bin when projected into the hierarchy measured at the original time point versus the hierarchy measured at P28. We find that the spatial patterns at the various time points agree significantly above chance (Figure 6**b**).

We also measure the correlation between the axes of covariance 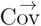 among the same hierarchical node at different time points, and among different hierarchical nodes at the same time point. We find that the axes of the same node across time correlate more than the axes of various nodes within a single time point (Figure S12).

### Hierarchy exists in small gene sets

As we have selected genes without bias, the spatio-temporal patterning may be a general property of gene expression, rather than the specific property of a particular set of specialized genes. We explored this possibility by randomly selecting sets of genes of various sizes, and then measuring how well the spatial patterns were maintained.

From the spatial gene expression data we found that small sets (*∼*50) of genes selected randomly from the database can already achieve an accuracy (ratio of voxels sorted in the same hierarchical bin) with the signal over all genes (Figure 7). So, the lineage identity of a cell may be encoded in the profile of any small set of genes.

**Figure 7:**
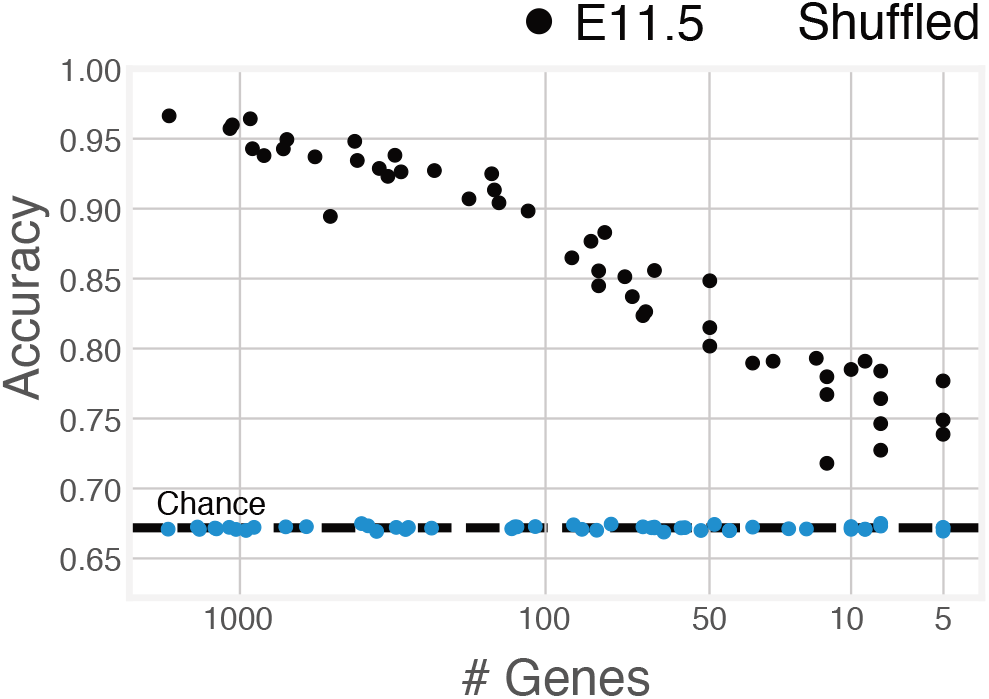
Random sets of genes of various sizes from embryonic age E11.5 were selected, and the spatial hierarchy they exhibit was compared to the hierarchy exhibited by the grand set of all genes at hierarchy. To compare hierarchies all voxels are projected onto both hierarchies. For each matching choice the score is incremented proportionally to the depth of the bin. As such, 1 indicates that the all voxels are sorted into corresponding nodes of the hierarchies, and the dotted line indicates the score if all voxels were sorted into hierarchical bins randomly (as in the shuffled case). The hierarchy established from a set of 20 random genes already agrees largely above chance with the original pattern.

### Simulated mitosis induces spatial hierarchy

We verified that our division model is able to explain the above results by simulating the mitotic development of a cell mass from a small pool of precursors (see Methods for a more detailed description of the simulation).

The simulation embodies the constraints of the model. The basic element of the simulation is a space-occupying (not a point) cell that expresses a profile of genes. When a cell divides, the expression profiles of its two daughters are normally distributed variants of the expression of their mother. The cells are additionaly positioned in a spatial 3D grid. When a cell divides it pushes a nearby cells in a random direction to make space to place the mitotic daughters adjacent to one another (see Methods). In this way, we are able to efficiently simulate the growth of a volume of cells, analogous to the embryonic brain. A particular lineage tree of cells is generated by recursive application of this division rule.

The simulation results in a mass of 500,000 cells, each expressing 500 genes, and distributed over 100 independent lineages. This mass was divided into 21140 voxels of 3×3×3 cells (excepting voxels on the outer surface, which may contain fewer cells), to emulate the voxelation of the ABI data. We then applied the same hierarchical decomposition as was applied to the experimental data, yielding qualitatively similar results (Figure 8). This simulation indicates that the spatial and genetic constraints of the model are sufficient to explain the spatial patterns that we observed in the developing mouse brain.

**Figure 8:**
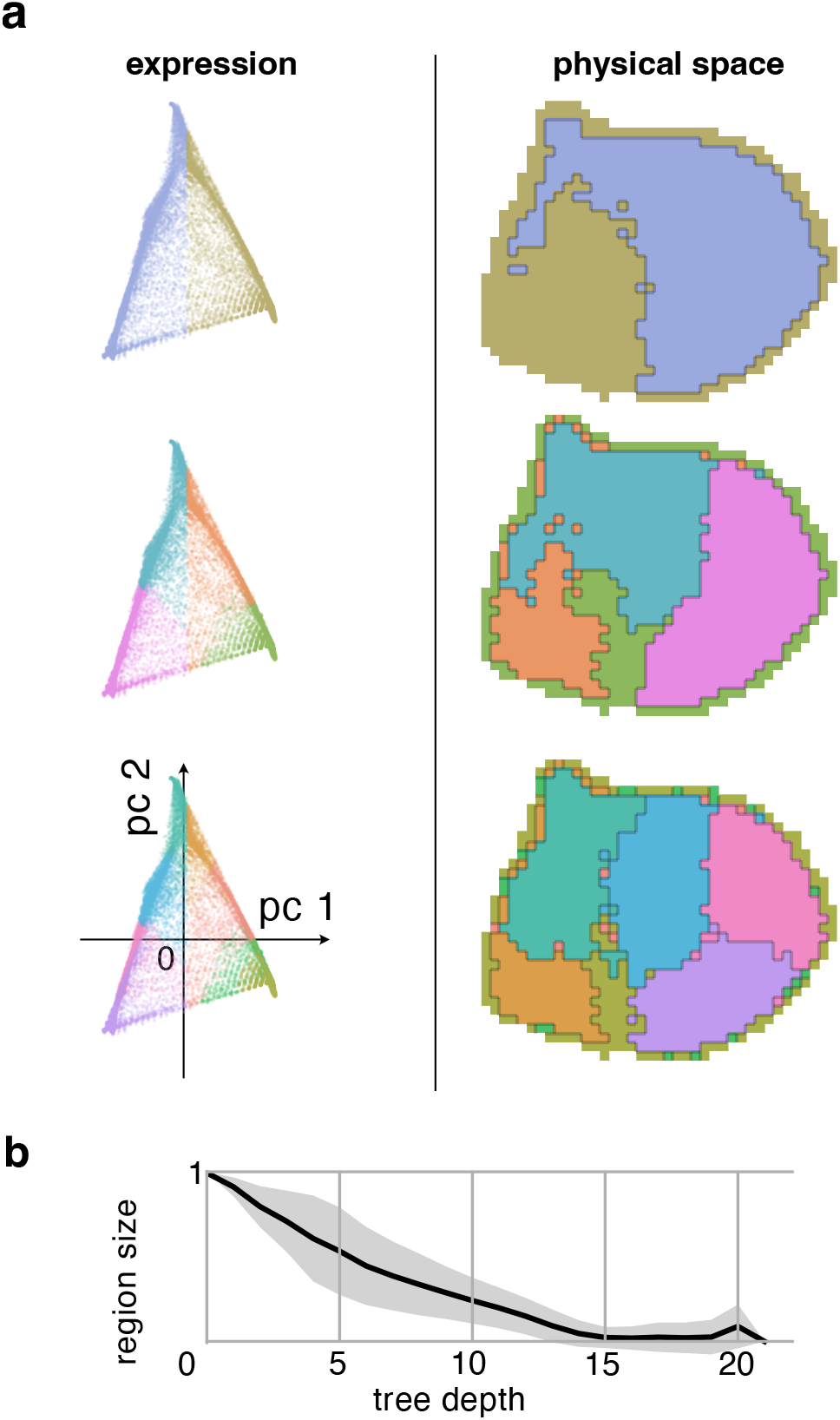
Hierarchical decomposition of expression data generated by simulation of the model (see text) proposed to explain the results. Simulated ‘brain’ sphere composed of voxelated leaf cells was generated by 300,000 mitoses distributed over 10 independent lineages. Cells express 500 genes. Asymmetrical mitoses induce differential changes in gene expression. Each voxel contains 3 × 3 × 3 = 27 adjacent cells. Similar to experimental results, the hierarchical decomposition of covariance in gene expression voxels independent of location (left), is mirrored by matching decomposition in space (right).

### Simulated axons traverse the brain

We simulate the process of axonal growth to demonstrate that they can use the Familial Guidance Model described in the Concept to navigate through the voxels of the ABI gene expression atlas (Figure 9). The Guidance Model in the Rationale was described in terms of individual cells and a given lineage tree. In applying that model to the experimentally observed ABI data, we have to consider that the expression data is voxelated, with each voxel containing many cells; the lineage tree is only estimated, as described in the previous section; and we cannot yet predict which exact set of genes participates in the address space. So, we need to test whether robust long range guidance is possible at all using our proposed mechanism, rather than predict specific projections.

**Figure 9:**
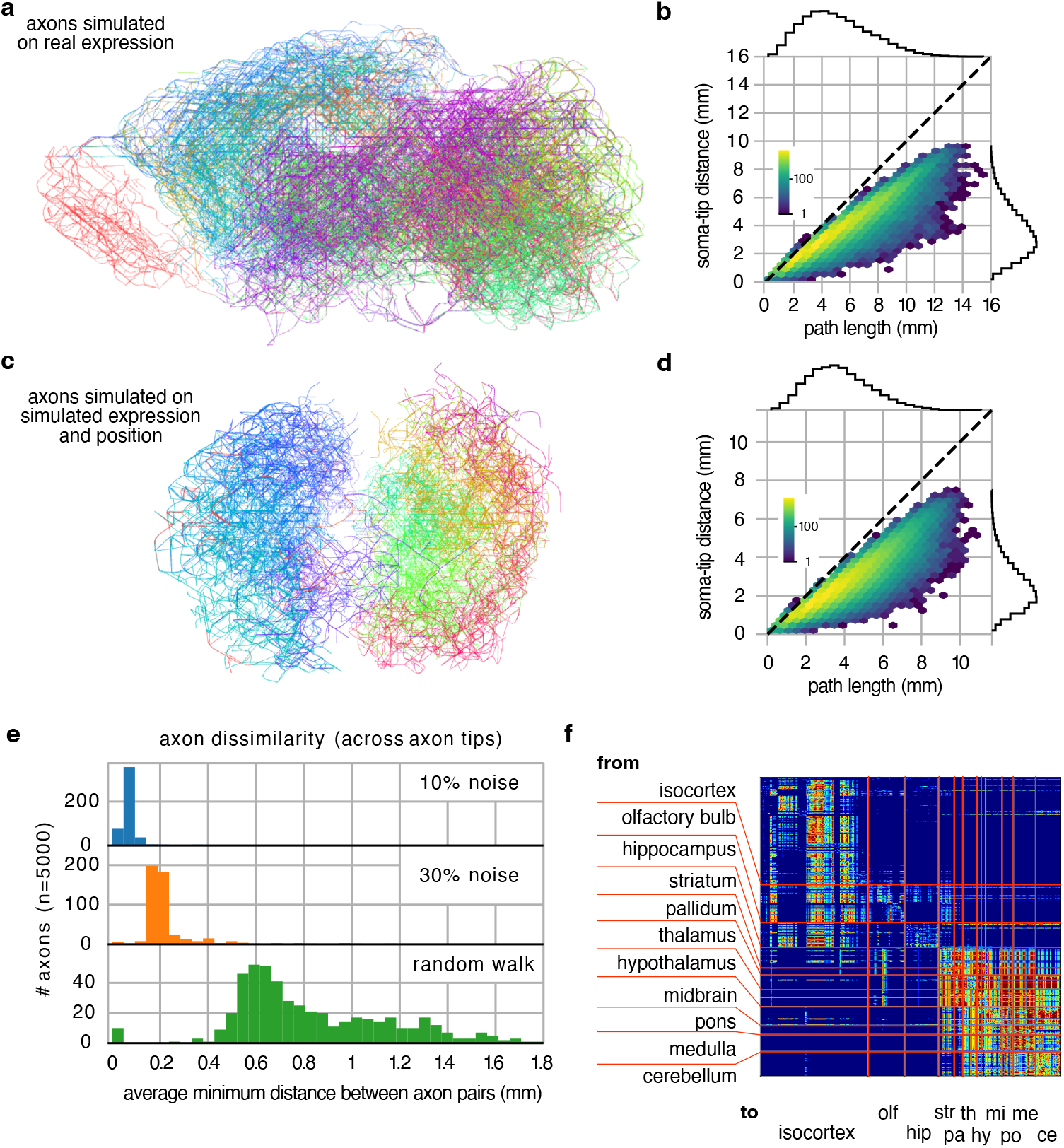
Simulated axons use familial guidance to navigate through the voxels of the ABI Developing Mouse Brain gene expression atlas. **a** Arborizations of 50 example axons, show in a sagittal projection of the ABI atlas. Each arbor is the collection of all branches that an axon could potentially navigate using this gene expression space. Each axon is colored according to its source region. The colorings correspond to those of Figure 5**b. b** Straight-line distance between the beginning of a branch (soma) and end of that branch (top) versus the actual path length. Branches are points sorted in hexagonal 2D bins, whose color intensity indicates the number of branches in that bin. **c** Same as **a** but on a tissue grown in simulation (as in Figure 8). **d** As **b**, but for the simulated tissue of **c. e** The dissimilarity between axons beginning from the same voxel (measured as average minimum distance), under varying levels (10% or 30%) of expression noise. (Because the navigation algorithm is deterministic the 0% noise case produces identical neurons.) The familial guidance dissimilarity is compared against a random walk axon of the same path length. **f** Connectivity matrix corresponding to the connections made by the axons of **a**. The connections conform to reasonable anatomical patterns. The anatomical regions marked on the matrix are taken from the annotations of the Allen Brain Institute. They are not used for the analysis.

The growing axons were simulated using a spatial-state graph approach. In this approach, the axon traverses a graph where each nodes represents the growth cone at a spatial location (i.e. a voxel) with a specific receptor configuration. Two nodes are connected by an edge if they are spatially adjacent and represent the same receptor configuration—this represents a move of the growth cone in space; or if they represent the same spatial location, and the receptor configurations are adjacent in the lineage tree—this represents a transition of the growth cone in state.

The axon can only traverse an edge to an adjacent node if the gene expression in that node is a better fit to its receptors than the current node (i.e. if the currently held ancestral state is more correlated to the adjacent node). This ‘biological’ algorithm resembles Dijkstra’s algorithm for finding the shortest route between two nodes of a graph [10].

Simulated axons were found to be able to extend up to about 10 µm source-tip distance, with axonal lengths of up to 16 µm (Figure 9**b**). The axon length is longer than the source-tip covered distance because the axons do not move in a straight line. Nevertheless, the relation between path and straight distance is close to linear for many axons, particularly those of shorter ranges.

The axonal guidance is robust against signal noise. We demonstrated this by adding noise to the gene expression after the lineage tree was reconstructed, but before the axons began guidance, effectively reducing the signal to noise ratio. The added noise does not significantly change the trajectories of the axons (Figure 9**e**). However, when gene expression is shuffled completely, either before or after reconstructing the lineage tree, axons fail to navigate beyond 1 or 2 voxels (not shown).

Similar axon length distributions are found from the fully simulated tissue (Figure 9**d**). This result further supports the hypothesis that simple constraints on mitosis can induce the address space.

The control case for axonal arborization is a random walk axon of identical total length, whose growth cones are still self-avoiding, but able to move in random directions at every step. We find that the random walk axons have much lower specificity, robustness, and spatial reach, than the navigating axons (See Figure 9**e**). Note that for the navigation algorithm the total length of the axon is an implicit result, rather than a set parameter. Thus, the random walk axon already over-informed compared to the navigating axons.

We generated a typical connectivity matrix by simulating 500 axons rooted in voxels sampled uniformly from the available data, (Figure 9**f**). The matrix is sparse and block structured, with many off-diagonal components (rather than narrow diagonal band), indicating a specific, regionalized, connection pattern. Remarkably, these typical connections conform to reasonable anatomical patterns, as can be appreciated by comparing the block-structured axonal connections with anatomical regions taken from the independent annotations of the Allen Brain Institute (Figure 9**f**. We emphasize that these anatomical annotations are not used at any point during simulation or analysis.

## Discussion

The literature describing the progressive organization of vertebrate brains over embryonic and evolutionary time [42, 2] has emphasized local organizational processes and their dependence on a local landscape of molecular signals [37, 1]. However, that focus neglects the more global question of how the guidance landscapes are themselves established. Long-range migration of neurons, extension of their axons, and the formation of their many synaptic connections require a global orchestration of guidance cues [17, 50] at various spatial scales.

In this paper we have explored the hypothesis that the mitotic lineage tree, which is implied by the cellular gene regulatory network (GRN), is key to understanding the necessary global orchestration of molecular cues. The lineage tree induces a global guidance address space over the embryonic brain that is encoded in profiles of expression of multiple genes. Some part of the expression pattern of each cell includes the precursor signatures that encode its ancestral path down the lineage tree. The expression signatures of early ancestors are broadly spread across the present progeny, whereas the distribution of signatures of recent progenitors is more restricted. These systematic differences in location and scale of distribution of ancestral expression patterns supports the address space. Because the address reflects family lineage, we call it a Familial Address Space (Figure 3).

We further propose that the address space is navigable by axonal growth cones, which are able to grow to specific target addresses by matching local gene expression patterns to those of successive nodes of a lineage tree traversal. We call this process Familial Guidance (Figure 4). In other words, the expression of the brain, and the growth cone’s ability to exploit it, are dual consequences of the brains developmental process which both creates the Familial Address Space as a consequence of cellular differentiation, and then exploits that differentiation for active cellular organization including the formation of axonal connections.

Molecular labels were proposed by Sperry to explain how retinal axons select their targets in the tectum [31, 48]. However, it has been unclear in how unique, dynamic, and matching labels could be simultaneously presented by the tectum and recognized by axons from the retina [56]. Particularly, in these and other explanations of circuit formation [11, 46] it is unclear how the reproducible connectivity can be encoded within the genetic budget. Our proposed mechanism resolves this issue by showing that the lineage tree can efficiently install unique labels in target tissue, and that navigating axons can recognize them due to their shared origin in the cellular GRN. It also extends the scope of comprehensive molecular labels from the retino-tectal projection to the brain at large.

We searched for evidence of such an address space in the ABI mouse brain atlases, because they provide voxelated (rather than tied to pre-conceived anatomical regions) spatial expression data of developmentally relevant genes throughout brain development. Previous analyses of these and related atlases have been largely concerned with identifying profiles of co-expression that support anatomical organization [37, 39, 38, 34, 51]; and also whether these regional profiles can be explained in terms of the known expression of specific cell types [18, 58, 60]. Although there are also systematic transcriptional similarities across cortical areas [34, 19], no global map-like organization has been reported as yet.

Our results now indicate that systematic spatial patterns of gene expression covariance do exist and are widespread in the embryonic and postnatal brain. These patterns involve non-specific groups of genes, occur on multiple spatial scales across the entire brain and spinal cord, transcend neuroanatomical boundaries, and are consistent at least from E11.5 to P56. Interestingly, we found that the primary axis of variance corresponds spatially to the dorso-ventral axis of the embryonic brain, rather than the antero-posterior axis that is expected on the naive assumption of greater variance along a longer axis. This suggests that the patterns do not simply reflect the geometry of the developing embryo, but are related to controlled regionalization in embryogenesis itself.

We explored the embryological origin of these patterns by analyzing the statistical structure of the expression covariance [20], rather than the relationship between expression and anatomy [34] or to phenotypic expression of cells [18]. The essence of this structure is that the differential gene expression between arbitrary sibling branches of a lineage tree (the asymmetry) in expression space has a dual expression as covariance across the region of brain space occupied by the leaf nodes of those sibling branches, as proposed in the Rationale section. Indeed, simulations of the Familial Address Space model showed good qualitative agreement with the experimental data (Figure 8). They confirm that the differential gene expression profiles induced by early divisions can be reconstructed from the gene expressions observed in the leaf cells of the lineage tree.

The covariance patterns indicate only that common gradients of expression exist across sets of genes, and seem to be hierarchically organized. Our results do not of themselves indicate which genes contribute most strongly to the patterns, nor which, if any, are actually utilized for addressing. It remains to be tested whether the spatial organization observed in the current data is restricted only to the *∼*2000 genes that the ABI chose for assaying [51], and consequently to the subset of *∼*1, 240 that we have analyzed. However, both the experimental data and our simulations indicate that the organization does not arise from the expressions of a specific set of genes dedicated to encode spatial structure, but rather can be found in the expressions of any sufficiently large (*> ∼*50) set of genes. Thus, the spatial hierarchy in gene expression depends primarily on the statistics of the induced changes, while the specifics of gene function are less relevant to their generation.

The range of spatial scales (Figure 5), temporal stability (Figure S11), and near orthogonality (Figure S12) of the covariance nesting is suggestive of an address space. This putative address space has an interesting property: Because the nested regions are a projection of the lineage tree onto 3D brain space, regions composed of cells that are closely related in their respective lineage trees are also close in space. Thus, the address map is a systematic arrangement of cells in terms of their ancestral gene-expression, and so provides an implicit encoding of cell lineages that could be used as a relative localization mechanism that can guide tissue organization. For example, migrating cells or individual axonal growth cones could steer to their target locations by tracing a sequence of address patterns. These patterns need not have evolved with the intention to guide growth cones: It is sufficient that a growth cone recognize the pattern and exploit it as a directional cue. Growth cones might recognize these patterns, because they are not only the product of the global developmental program, but themselves contain that full program in the genetic code of their source cell. Thus, we may expect migrating cells and cellular excrescences such as axons to have methods of decoding that and relate to mechanisms by which the expression patterns are themselves induced.

While constructing the address space, the mitotic tree is rooted in the stem state of its gene regulatory machinery. However, if the leaf cells root their regulation in their own current states, then their potential exploration paths are traversals of regulatory paths to destination states, as seen from their origin state. Thus, the exploration paths of the growth cone can be seen as the lineage tree hung from a leaf (with some pruning). So, growth cone routes are anti-differentiating up the tree to some ancestral node, followed by re-differentiating toward the leaf states accessible from that ancestral node.

An appealing aspect of this lineage-induced address is that it greatly simplifies the evolution of complex spatial organization of cells. The systematic spatial labeling of cells is given as a direct consequence of mitotic specialization and cell proximity. Evolution needed only to discover how to exploit this labeling for organizational migration of cells and their components (e.g. growth cones). It could opportunistically select a set of gene products for axonal growth cone guidance, because most gene sets will encode a similar spatial pattern. This generality of the address space could also help to explain the wide range of guidance cues that have been documented [50, 28, 45]. The selection of a subset of cell surface markers, or diffusible markers would be a convenient choice for growth-cone sensors.

The Familial Address Space model is entangled by two factors. Firstly, the mitotic root of the developing brain is difficult to define exactly. It seems reasonable to consider as starting point a small collection of early progenitor cells downstream from the zygote that are committed to formation of the neural tube, rather than the single progenitor that we have assumed for simplicity in describing the model. Secondly, the experimental data are voxelated and so average over the various cell types (possibly derived from different lineage trees) that they contain. These two factors will mix and average the effects of the simple model. Nevertheless, if the lineage of mitoses is sufficiently coherent in time and brain space, then the statistical signature of the mitotic process remains detectable. The spatial patterns persist even when confounding mechanisms, such as symmetric cell division, de-synchronized mitotic clocks, data voxelation, and multiple independent lineage trees are introduced to the simulation (Figure 8).

Gene expression is a central aspect of our Familial Address Space model. The gene expression of a cell is a 2000-element vector, which encodes the expression energies of the *∼*2000 genes used by the ABI atlas. Since the exact expression profile of the root cell is unknown, we assign to it a fixed pseudo-random number. The profiles of its progeny are obtained by successive applications of *5*s, drawn also from a frozen generator. These frozen stochastic expression profiles and their transformations are merely a convenient proxy for the unknown (deterministic) sequences of gene expression over consecutive cell division that occur in individual cells during development. The actual sequences of expression are not crucial to the model because it is the induction by mitosis and then the propagation of the statistical signal that is of interest here, rather than the absolute expressions of particular genes. We may also allow that the stochastic profiles be subject to cell-external or internal factors, provided that these influences are reproducibly regulated as part of the developmental process (and thus not due to environmental noise external to the embryo).

### Axonal connections by Familial Guidance

There has been substantial progress in understanding how axonal growth cones respond to local guidance cues [5]. They are exquisitely sensitive to local gradients, able to detect gradients on the order of a few molecules across their span [41]. However, physical noise, ligand binding, and other signal detection considerations indicate that molecular gradients alone are insufficiently robust to explain axonal guidance, particularly at longer spatial scales [17, 5]. Over these longer distances the algorithmic rather than the reactive aspects of guidance rather relevant.

Previous models have described network formation in terms of cellular agents containing a small, but explicit, program consisting of few developmental primitives [63, 64, 65, 4, 3]. These generative cellular programs include physical constraints on development to explain network formation [21]. Our work puts such generative algorithms in a broader context, by showing how physical constraints induce an address map essentially without any explicit program. The generation of the address map acts as an organizing principle that more specialized cellular programs might exploit. Here, we have shown that even a very generic axonal algorithm, Familial Guidance, is able to install a basic connectome.

The Familial Guidance Model is cast as a growth cone guidance algorithm. The algorithm depends on the embedding of the lineage tree in both expression space and brain space (Figure 4). The axon navigates by entering a control loop that first uses the inverted developmental program to revisit an ancestral expression state, and then configures the receptors in the growth cone to search for marks of that expression in the surrounding tissue. When the local optimum is reached, the axon transitions to another ancestral state and the process repeats, until another leaf is found. The growth cone’s choice (i.e. reconfiguration) points reflect transitions through the ancestral lineage tree: The growth cone knows how to reconfigure its receptors appropriately because it is able traverse (in expression space) the lineage tree.

To find its target an axon must trace a route through physical brain space. Growth cones at the tips of axonal branches guide the extension of axons by receptors in their membranes. These receptors recognize morphogenic cues [50], and also membrane bound makers [28], and either promote or prevent the extension of the axon in the direction of increasing cue concentration or prevalence. The cones dynamically change the profile of receptors in their membrane so as to change the criteria for direction sensing at discrete way-points [45]. In our model, these growth cone receptor profiles correspond to expression signatures (‘marks’) of ancestral cells. That is, the receptors recognize these ‘marks’ in current cells that they have acquired by virtue of being the progeny of those ancestors. A guidepost cell would then be an early born cell, in which these marks are still strongly expressed. Figure 3 explains how expression covariance patterns are induced at cellular level as δ changes in *c*_*i*_, but that the global address space arise is observed over whole populations of cells as 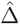. And in this paper we have emphasized the experimental observations of lineage address space composed of the ordered. 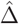. But, of course, to make use of this lineage address space for guidance, individual grow cones will have access only to the local *c*_*i*_, and their resultant guidance cues secreted into the extracellular space or exposed on cellular surfaces.

The model asserts that the navigation sequence can be generated if the axon inverts its developmental program to anti-differentiate to precursor expression states, and thereby traces a route through the lineage tree. There is evidence that cells and also neurons are able to de-differentiate as a whole [14, 30, 57]. However, we require only that de-differentiation occur on some subset of the genome relevant the familial marks. While there is as yet no systematic work on this question, there is nevertheless clear and growing evidence that growth-cones used elaborate local context-dependent mRNA processing during their guidance behavior [55, 25, 9].

Our guidance algorithm requires that axons perform a virtual traversal through the neuronal lineage tree to generate a physical route map. This conformity is possible only if (1) the tissue retains a persistent record of the lineage tree, so that expression signatures of ancestral cells can be recognized in their progeny after the ancestors have vanished (through mitosis), and (2) routes through the lineage tree are indeed matched (at least in relevant cases) by continuous routes of adjacent progenies through the tissue. We found evidence for these two conditions in voxelated gene expression data. The forward projection of a hierarchy measured at an early embryonic age matches that measured at later age (Figure 6, S11) indicating that the gene expression of cells holds a persistent record of their ancestral gene expression profiles. And secondly, the voxels of the grand lineage tree estimated without regard for location, are nevertheless grouped together in space at each tier of the hierarchy so forming adjacent and contiguous regions (Figure 5, S13–S20), indicating that lineage paths may indeed form continuous trajectories in the tissue.

We tested the Familial Guidance algorithm in the original voxelated data. Because we do not have access to the true mouse lineage tree and its genetic states, we used the reconstructed lineage tree obtained by hierarchical decomposition of the voxelated data as an estimate the growth cone reconfiguration transitions. The growing axons were simulated using a spatial-state graph approach. These simulations confirmed that axons do indeed grow to more specific and longer range targets than the random walk model, and that an arbitrary collection of axons reproduce qualitatively the sparse and block structured connection matrix of the kind observed in experimentally observed connectomes. These typical connections conform to reasonable anatomical patterns, as can be appreciated by comparing the block-structured axonal connections with anatomical regions taken from the independent annotations of the Allen Brain Institute (Figure 9**f**. Although this general agreement is in itself remarkable, the particular connectome is not yet a proper prediction of actual connectivity. Several issues will need to be resolved in order to improve the prediction. Obviously, the range and resolution of experimental data must be improved: the ABI data offers only subset of genes, and even these data degraded by averaging over voxels and brains. Furthermore, the address space was inferred from the set of all genes. However, this set is only one of many possible sets of genes that support the true address space (Figure 7). It is likely that evolution has selected a particular set of genes to establish a particular address space that supports well evolution’s preferred connectivity. Unfortunately, this particular set of genes is as yet unknown to us, although there are some strong candidates for inclusion [24]. Furthermore, the simulations themselves are restricted. Guidance was only simulated at one time point, consequently the trajectories are generated as if development were frozen in the P28 geometry and gene expression. Of course, other trajectories will be possible at different developmental times. And for reasons of computational resource, sampled axon sources were a randomly sampled subset of all voxels, so many trajectories are omitted: It would take prohibitively long to simulate all sources.

Overall though, the address space induced by mitosis, as well as the guidance process that it supports, is consistent with reasonable axonal projection pattern. Full agreement between our simulations and experimentally observed projections will depend on the agreement between our estimated differential gene expression model, and the true differential gene expression generated by the actual gene regulation network of the mouse. These simulations also confirm at least a partial projection of the expression space onto brain space. This conformity is not self-evident, because the high dimensional expression space cannot be faithfully projected into the lower, three dimensional brain-space. Even in the best embedding, the pairwise distances between the embedded nodes in 3D Euclidean space cannot exactly match the pairwise path distances between nodes in the tree. The error can be made arbitrarily small in the limit of many dimensions. This limit is not relevant for 3D physical space because its dimensionality is fixed; but for gene expression space it is relevant because the dimension can be increased by recruiting genes for the embedding. Fortunately, constraints on mitotic daughter migration will result in at least some regions of continuity in the lineage tree embedding. Thus, although not all traversals through the lineage tree will be matched by traceable paths in brain space, those traversals whose embedding in brains space provides for continuity of expression signals will be successful. This property is reflected in that our connectivity matrix is not fully connected or diagonally structured, but sparse and block-structured (Figure 9**f**). The manifold of traceable paths, and so connection probabilities, will no doubt be influenced by the anatomical distortions imposed on the growing cell mass by factors such as relative mitotic rates, cell size, asynchronous axonal outgrowth, ventricular volumes etc.

Note that our algorithmic approach differs from more usual methods for the generation connections, such as a connection table of source destination pairs, or a graph generation rule (e.g. Erdős–Rényi) that connects nodes according to a statistical model. For example, a common connectome generating rule is that Euclidean distance between pairs of neurons be inversely proportional to their connection probability. In this case, two nearby neurons are more likely to connect than distant neurons [13]. A typical implementation of this rule would involve measuring the Euclidean distance between two neurons, and then deciding whether a connection exists between them by evaluating a probability distribution. This method will establish suitable entries in a table, but does not explain how these connections should be grown in space. To satisfy the rule by a developmental algorithm the growth cones must perform a random walk in space, governed by a fixed probability of extension; and connect to encountered neurons also with some fixed probability. This simple connection algorithm closely reproduces the empirically observed axonal length distributions. However, because the behavior of these parameterized stochastic models depend on random data (rather than fixed data), they are also unable to explain the repeatability (across individuals) patterns of axonal trajectories and connectivity observed in biology. Repeatability would require that the ‘random’ walk be decided by a frozen random number generator, so simulating a deterministic guidance mechanism, whose data is that frozen random number.

A traversal of axons through the lineage tree explains the experimental finding that cortical excitatory cells seem to preferentially target their clonal siblings [8, 27, 59], rather than simply nearby targets.

Our axonal growth simulations are only for pioneer axons. Many other axons may reach their remote target through fasciculation with a pioneer axon than by pure pioneering themselves [47]. However, these follower axons could still use the same guidance mechanism as pioneers to make decisions for (de-)fasciculation, so avoiding additional encoding as to when to fasciculate with which other axon. (If such a fasciculation specific route encoding were necessary, it would probably require on the same order of information as the naive wiring diagram, depending on how prominent fasciculation is.) An elegant solution would be that each axonal segment maintains the expression state of the growth cone when it created that segment. In this case the growth cone is seen as any other axonal segment, except that the growth cone is motile. In this way the axon segments become strong markers, expressing the signal that other growth cones can follow. Their signals would be exceptionally strong because the growth cone imparts to each segment the ground-truth ancestral signature obtained from source genetic information, rather than the a noisy signature that has been projected through generations of progeny. This address efficiency could explain the observation that the growth cone of a fasciculated axon is only a fraction of the size of a pioneering axon’s growth cone. Such an Ariadne mechanism would permit late growing axons to traverse areas whose geometric continuity with the lineage tree existed earlier during development, but has since been disrupted.

### Genetic encoding of Familial Guidance

The genetic (and epigentic) information required for instantiating the physical neuronal network is largely limited by the roughly 1GB information capacity of the original zygote. Evidently, the detailed physical network arise through a decompression of this information, resulting in the connections summarized by the connection matrix. A naive encoding of this matrix for mouse brain connectome would require roughly 10TB. However, viewing the connectome as a generic connection matrix considers too many possible configurations, and consequently overestimates the information necessary to specify the brain’s connectome among them. No doubt there are regularities in axonal construction that can express various arborization types using simple codes [64, 22], and there are means to generate connectomes from compressed codes that do not suffer from the constraints of the construction process [26]. However, the many disparate long range pathways of the brain would require more elaborate codes and co-ordinate systems. The Familial Guidance principle shows how the implicit, compressed, representation of the target connectome can be decompressed through the very construction of the neurons to be connected. Self-replication, with its inherent constraints, organizes the growing mass of cells in a family hierarchy whose parent-child relations manifest as spatial gradients of differential expression. These gradients act as a network of roads that axons explore to reach their targets. The directions for axons to establish a default wiring in this familial landscape are simple: grow and branch along every accessible road. The final network and the landscape are hand in glove. In this view, growing the landscape is at least as relevant as axonal outgrowth. But fortunately the growth of the fundamental landscape follows very directly from simple constraints mitosis, and so has low informational cost. In this view, the 1GB genome contains no compressed 10TB connectomic blueprint for the brain. Rather, the genome encodes the host cell, which is essentially a self-replicating physical machine [33] whose execution (or decompression) generates the wired brain. Therefore, the size of the brain’s blueprint is not limited by the size of the genome, just as the uncompressed size of the output of a computer program is not limited by the size of its source code.

## Conclusion

Our analysis of gene expression in embryonic and postnatal mouse brains reveals a hierarchy of spatial patterns of expression covariance that extend over the entire brain, and are stable over the available data. This organization is present across the 1240 genes analyzed. However, they are also present in the expressions of random subsets of as few as 50 genes. The organization is consistent with a multi-scale address space that could be exploited for cell migration or growth cone guidance. Our simulation studies confirm that this organization can be generated by persistent asymmetries of gene expression introduced by the successive mitoses of the lineage trees that give rise to the brain, provided that mitotic daughters do not stray too far from one another after their birth.

Due to the generality of these mitotic constraints, it is likely that similar map-like structures exist also in other tissues, and may provide a fundamental scaffold for cell migration and tissue organization. However, the Familial Address Map has particular relevance for neurons, whose many stereotyped connections cover distances up to the scale of the whole brain.

We conclude that the fundamental wiring of brain can be compactly encoded and expressed through the mitotic lineage implied by the genetic code of its embryonic stem cells, because the arborizations of axons are just the available search paths through lineage tree. So, paradoxically, (cell) division may be the key to uniting the neurons of the brain. The resolution of the paradox is that division in reverse is unification.

Future work must establish: which specific sub-set of genes is used for axon navigation; how the growth cone reverts its host’s differentiation and how receptors are generated to recognize an ancestral state; and how the address space, that is the geometry of the brain and spatial gene expression, are tuned to realize a specific observed connectivity.

Contrary to the prevailing reductive approaches to understanding the wiring of the brain, this paper has taken a more global synthetic view. While much more effort will be required to confirm the various implications of our approach, the theory and available data are remarkably consistent; and offer the prospect that the connectome and its functioning can be more readily understood in terms of the global mechanisms that generate it, rather than from interpretation of the final wiring diagram, just as inspecting source code is more revealing of principles of operation than inspecting the compiled program.

## Data availability

All experimental data analyzed in this paper were originally published by the Allen Institute for Brain Science [51], and are available at https://developingmouse.brain-map.org.

## Code availability

All custom code is available at …(will be added with paper publication)

## Acknowledgements

The authors acknowledge the Allen Institute for Brain Science for their mission to bring systematic neurobiological datasets to the public domain. Without their work, the research reported here would not have been possible. The authors also thank, Bruno Averbeck, Florian Engert, Richard Hahnloser, Denis Jabaudon, Henry Kennedy, Sepp Kollmorgen, Kevan Martin, David Price, and Saray Soldado for discussions and comments regarding an earlier version of this manuscript. This work was supported by ETH–24 19–02. The authors also thank Prof. Dr. Tobias Delbrück for support.

## Author Contributions

Data acquisition SK, GM; Analysis of data SK, GM; Simulations SK; Theory SK, RD; Wrote the paper RD, SK, GM; All authors revised and edited the completed document; Proposed the study RD, GM.

## Methods

### Experimental Data

The analyzed gene expression data were published by the ABI in their Developing Mouse Brain Atlas [51]. The data are provided as 3D grids of isotropic voxels of various sizes. The expression energies of the *∼*2000 genes were measured by *in situ* hybridization and take any non-negative value, while -1 indicates an invalid measurement in that voxel. ‘Expression energy’ is a combined measure of density and intensity. The voxel dimensions are 80 µm, 100 µm, 120 µm, 140 µm and 160 µm for developmental time points E11.5, E13.5, E15.5, E18.5, and P4, respectively, and at later time points, i.e. P14, P28 and P56, the voxel dimension remains constant at 200 µm [51]. Every voxel thus contains the cumulative expression of many (probably thousands) cells.

The atlas data was retrieved through the API provided by ABI. The ABI expression grids were used as published, without performing any additional re-sampling or interpolation (see below for preprocessing). Thus, the voxel sizes were maintained as published by the ABI.

Only measurements from sagittal sections that were not labeled as failed images were used (omitting failed and coronal sections). When multiple successful experiments were available for a particular gene at a particular time point, one of the experiments was selected arbitrarily.

From the 3D expression grids, only those voxels labeled (by the ABI) as part of the neural plate were selected. This includes all developmental derivatives of the neural plate, i.e. voxels of brain and spinal cord tissue, but omits those of ventricles and empty space. All individual voxels that have more than 20% invalid measurements and all genes that have more than 20% invalid values across all remaining voxels at every of the developmental time points (in that order) were removed from the analysis. In this way, the same set of 1240 genes was selected for each of the time points. The number of selected voxels are 7377, 12266, 11869, 11639, 21348, 24224, 28476, and 60129, for the time points E11.5, E13.5, E15.5, E18.5, P4, P14, P28, and P56, respectively.

To avoid the introduction of spatial confounds, the ABI recommendation to spatially interpolate remaining invalid expression values was not applied. Instead, missing values were replaced with the mean expression value of that gene over all voxels at that developmental time. Thus, when the data is later centered for analysis the invalid expression values become 0.

In order to make the gene expression energy levels roughly comparable across genes, the expression values were normalized to unit variance and zero mean over the voxels at that developmental time.

### Hierarchical Decomposition

The voxels measured at one time point are sorted into the leaves of an estimated lineage tree through our hierarchical decomposition procedure. The procedure starts at the root of the tree, to which all voxels are initially assigned. Each iteration of the procedure evaluates the voxels assigned to a node of the tree, and reassigns each voxel to one of the node’s two daughters. The procedure stops once every leaf node is assigned exactly a single voxel.

An iteration considers the gene expression of the voxels assigned to a node. The collected expression can be expressed as a matrix *X*, where each row corresponds to a voxel and each column to a gene. The voxels will be split over the daughter nodes along the axis of greatest variance.

The axis of greatest variance is the eigenvector corresponding to the greatest eigenvalue of the covariance matrix. The covariance matrix is computed by first centering the data by subtracting the empirical mean from each column 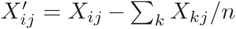, where *X*_*ij*_ is expression of the *j*th gene (column) in the *i*th voxel (row), and *n* is the total number of voxels (rows). The covariance matrix is then *Q* = *X*^*′T*^ *X*^*′*^.

The main axis of covariance is the eigenvector 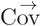 corresponding to the largest eigenvalue *λ* such that *Q* 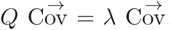. The eigenvector 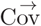 corresponds to the first principal component of gene expression covariance.

The coefficient *w*_*i*_ per voxel *i*, obtained by projecting the original data onto the axis of greatest covariance 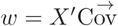 corresponds to the agreement of the voxel’s expression content with the axis 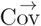. Based on these coefficient we sort the voxels into two subsets, namely one set (arbitrarily denoted *L* for left) with *L* = *{i*|*w*_*i*_ *<* 0*}*, and *R* = *{i*|*w*_*i*_ *>* 0*}*. These voxels of these sets are assigned to the left and right daughter nodes, respectively. The decomposition procedure is then repeated recursively on these two daughter nodes.

If a node is assigned only a single voxel, the process terminates for that branch. The process as a whole terminates when all branches have terminated.

### Controls

Controls were performed to ensure that the observed spatial patterns are due to the spatial distribution of experimental gene expression rather than being due to any inherent properties of the analyses. The null-hypothesis for the spatial patterning of expression covariance is that gene expression covariance is not spatially organized. In our control case, all gene expression values were permuted randomly across voxels and genes. This ensures that the overall statistics of gene expression remained identical, while removing all spatial structure from the source data. When the analytic workflow was applied to this synthetic data we obtained the results. These results confirm that the development of the mouse brain is associated with a systematic spatial organization of measured gene expression covariances, and that this organization is consistent over the period E11.5 through P56 (see Figures 6,S13-S20).

### Simulation of Mitotic Model

Our model of cell division was simulated numerically to confirm that constraints on gene regulation and mobility during cell division indeed induce a hierarchical gene expression address space. It also form one of the two substrates—next to the gene expression data grids from the ABI—for our simulations of axon navigation.

The division model has three components: a model of gene regulation that determines the expression profiles of the mitotic daughters at mitosis; a mitotic clock that initiates mitosis at some interval, and exits the cell cycle after some condition is met; and a rule for the placement of post-mitotic daughter cells in 3D space. In the minimal version of the model used here, a global clock initiates mitosis after an interval drawn from a Poisson distribution from the birth of the cell, and the daughters are born alongside one another along a randomly selected axis in the spatial simulation system described below.

A total of 200,000 mitoses distributed over 100 lineage trees, were simulated as follows. First, the topologies of 100 lineage trees were generated; next gene expressions were assigned to all cells; and finally the lineages were instantiated in model space. We chose this staged approach to the simulation for convenience of verification, and analysis. Simulations were written in the Python and C/C++, and run on a laptop computer. Code and documentation will be available upon publication.

#### A.4.1 Cellular Gene Expressions

Each model cell has a profile of gene expression, consisting of 500 genes. This profile is expressed mathematically as a vector of 500 values. For convenience, these values can be both positive and negative, which can be interpreted as positive or negative deviations from a base expression level.

Algorithm 1 describes the assignment of profiles in detail. In brief, the expression profile of each cell is a random variant of its parent’s expression profile.

Although randomness is used to establish the expression profiles, the (frozen) random deviations are used as deterministic process with statistics that are indistinguishable from a random process. This is analogous to fixing the seed of a random number generator: when the seed is fixed, the exact sequences of numbers is reproduced, but the statistics still seem random.

Of all cell divisions, 20% are symmetric (i.e. the gene expression of the daughter is equal to gene expression of the parent), and the others are asymmetric.

##### Algorithm 1: Gene expression

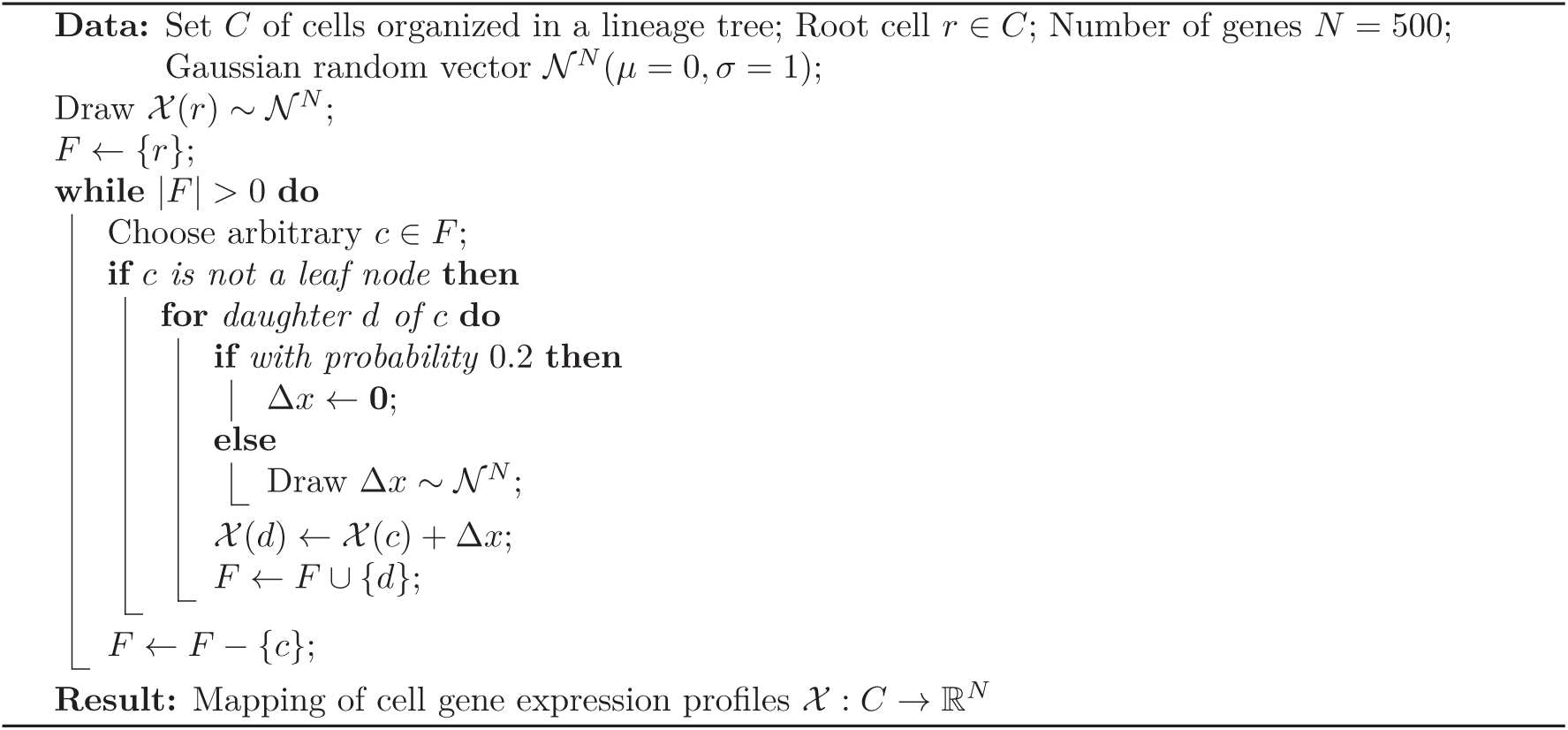

#### A.4.2 Mitotic clock

The mitotic clock mechanism generates the lineage trees by deciding when individual cells divide. It is described in Algorithm 2. When a cell is born, it draws a cell cycle duration from an exponential distribution (so that the process is Poisson). The division of that cell is then scheduled at the current global time, plus the drawn duration. At each iteration, the global timer progresses to the cell that divides next. The algorithm terminates when a fixed number of divisions is reached. Because the cycle durations are randomly drawn, resulting trees of varying number of nodes and with a generally unbalanced topology (i.e. branches have different sizes).

(The mitotic clock is thus independent of gene expression.)

To create multiple lineage trees, the algorithm is still performed only once, but starting not from one, but from multiple root nodes. So, the total number of divisions is possibly divided unequally over the lineage trees.

#### A.4.3 Cellular Locations

The cells of the lineage tree are positioned spatially by Algorithm 3, as illustrated in Figure 10. We developed this algorithm because it is simpler and more tractable than a direct simulation of soft (or solid) body physics.

**Figure 10:**
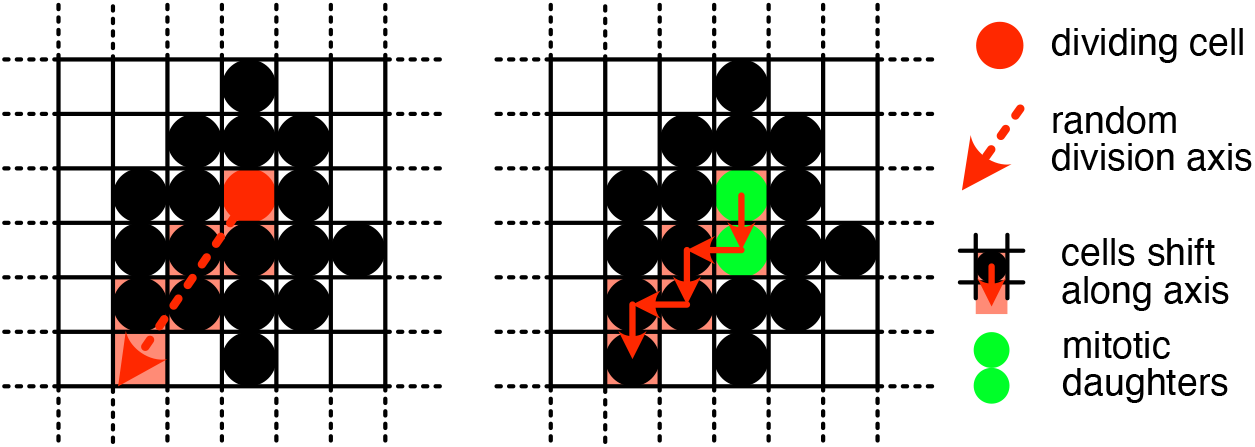
Illustration of cell placement. Although the illustration is 2D, the placement is the same for 3D. When a cell divides a random division axis is drawn uniformly from a unit circle (in 2D) or sphere (in 3D). Then, the sequence of cells that intersect the division axis are shifted along the sequence to create a free slot next to the dividing cell. The mitotic daughters take the original slot of the parent, and the newly created free slot.

##### Algorithm 2: Mitotic clock

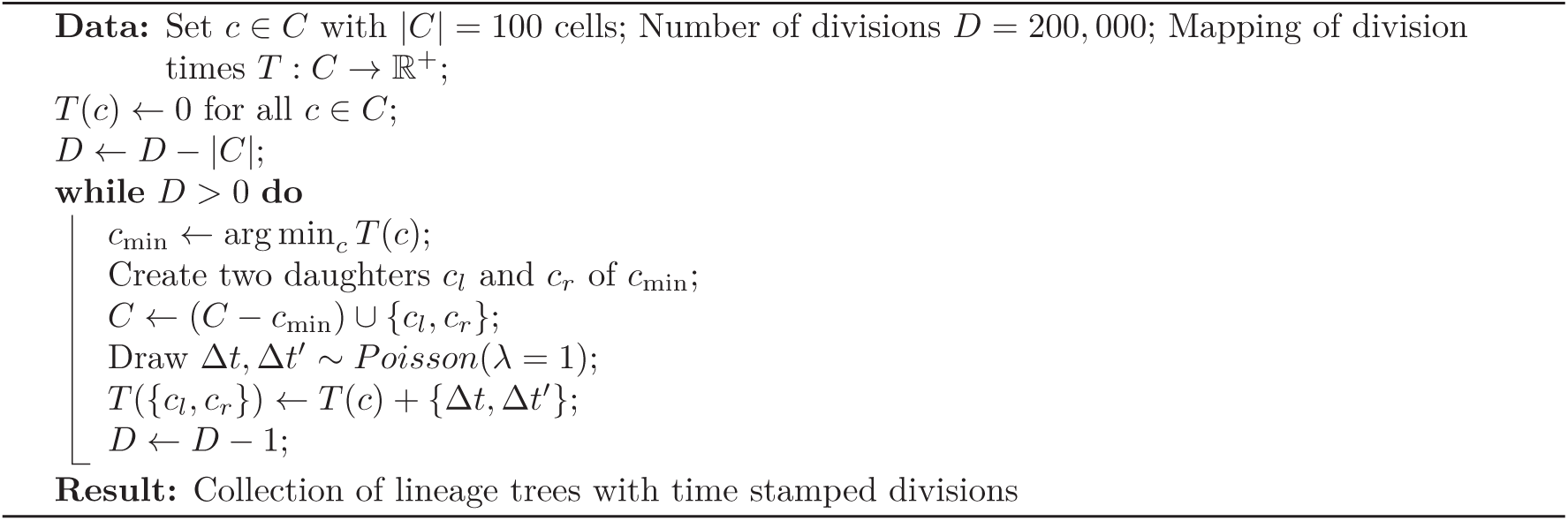

##### Algorithm 3: Tissue growth

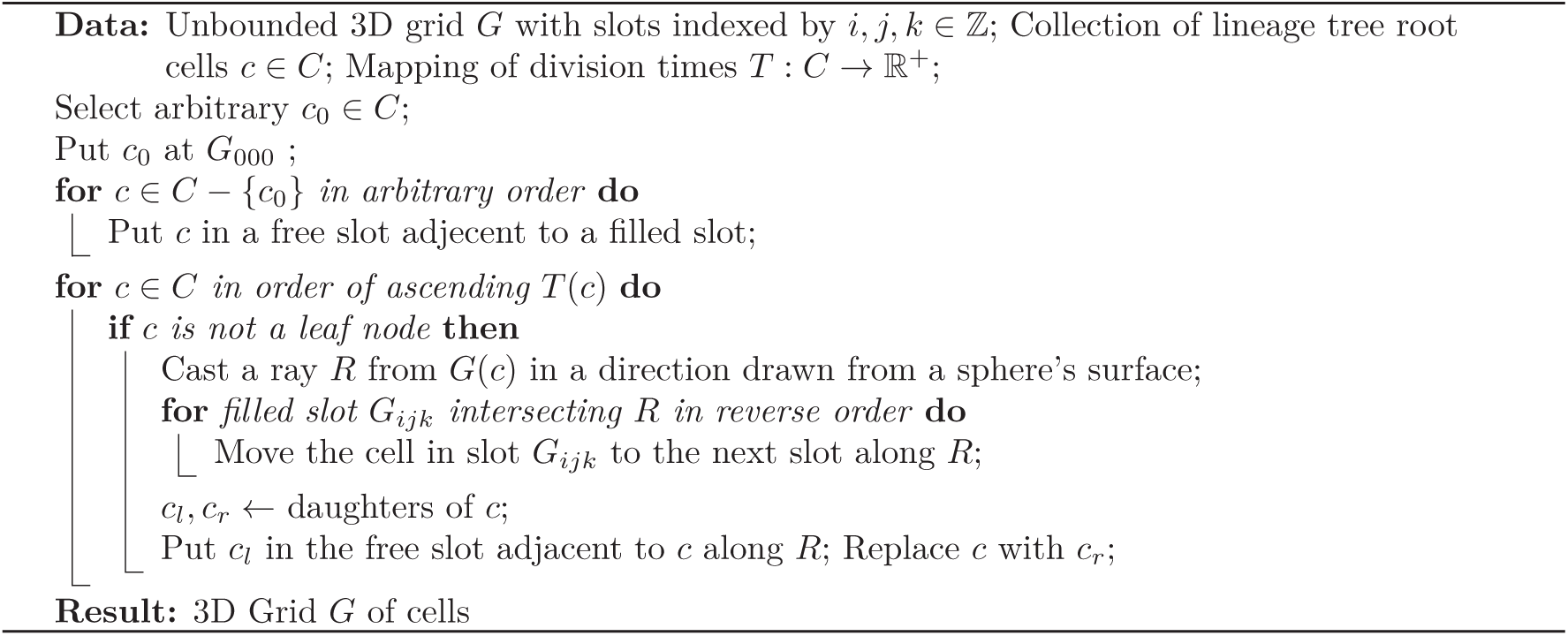

### Simulation of Axons

Our Familial Guidance model is simulated virtually, either on the voxel grid of measured expression, or on a grid of simulated cells.

The axon begins in some chosen leaf voxel, and takes as its initial template the expression state of its leaf voxel. Then, at each step, the growth cone senses the expression of the voxel it occupies, as well as expressions of the immediately adjacent voxels. The cone then moves into the adjacent voxel whose expression is most similar to its current template, so extending a new axonal segment between the traversed voxels. If multiple adjacent voxels contain a favorable expressions, then the growth cone is cloned, and the axon branches into all of those favorable voxels. Additionally, a cone can change (irreversibly) its present expression template to a that of an adjacent state up or down the ancestral lineage tree. The growth cone then repeats its search for favorable translations, on the basis of this new template. Each cone is constrained not to re-visit voxels already occupied by the cell’s axonal arbor (self-avoidance), and not to re-visit lineage states that it has already visited. The guidance process terminates when the growth cone can neither move to a more favorable adjacent voxel, nor change lineal state.

In executing this search algorithm, the initial cone and its clones extend axons along all the routes in brain-space that offer contiguity in brain-space of familial expression patterns encoded in the lineage tree (Figure 4).

For our axonal simulations brain space is discretized: Each spatial position corresponds to a (measured or simulated) voxel. Spatially adjacent voxels are connected by an edge. These nodes and edges form a graph encoding the geometry of the brain. The 3D positions of the nodes are used to establish the spatial graph, but ignored thereafter.

The adjacency of nodes is established through a Gabriel tessellation, which is a subgraph of the Delaunay tessellation [15]. In a Gabriel tessellation a edge of the Delaunay tessellation is kept only if the sphere of which the edge is the diameter contains no other points. This criterion ensures the spatial graph is connected, but that there are no edges across large empty spaces, such as ventricles and contours. This is an improvement over the vanilla Delaunay tessellation, which always contains the convex hull of the points, and therefore connects, for example, the rostral tip of the olfactory bulb to the cortex.

To navigate, axons follow signals on the spatial graph. A signal on a graph is a scalar value associated with each node, and the gradient of the signal is a value associated with an edge, that is the difference between the values of the nodes. The gradient depends on the direction the edge is traversed, and swaps sign if the edge is traversed in the reverse direction.

The signal for a growth cone depends on the current state of the growth cone, and the expression of the nodes of the graph. The state of the growth cone is a profile of gene expression that corresponds to a node in the lineage tree. The signal over the nodes of the spatial graph relative to a growth cone state is the correlation between the growth cone state’s expression profile, and each node’s expression.

For efficiency, the signal is only allowed to exist in the progeny of the ancestor whose state the growth cone has adopted. This constraint reduces the search space of the growth cone significantly, without significantly changing the routes taken by the growth cones.

The prominent action of the growth cone is to climb this signal by spatially moving across the graph, each time moving in the direction of positive gradient. If the signal value cannot be improved through moving, the growth cone has reached a (local) optimum.

In addition to moving, the growth cone can also change state. The state machine governing the transitions the growth cone can take is (isomorphic to) the lineage tree, which is estimated through our hierarchical decomposition. So, the growth cone can only transition to the parent state, or either of the daughter states, of its current state.

To simulate this process, a spatial-state graph is constructed. The nodes of the spatial-state graph are the Cartesian product of all spatial nodes, and all states. The nodes of this graph are connected if either the nodes are spatially adjecent, and have identical states, or if the nodes are spatially identical and have adjecent states in the lineage tree.

On the spatial-state graph there is only a single guidance signal, attributing to each node the correlation between the node state’s expression profile and node’s expression.

The navigation of an axon starting from a voxel is simulated by executing Dijkstra’s algorithm [10] from a source node in the spatial-state graph to all possible nodes containing leaf states, allowing only movements along positive gradients. For graph implementations the igraph library with python bindings was used (https://igraph.org).

Axons were visualized using threejs (https://threejs.org)

## Supplementary Figures

**Figure S11:**
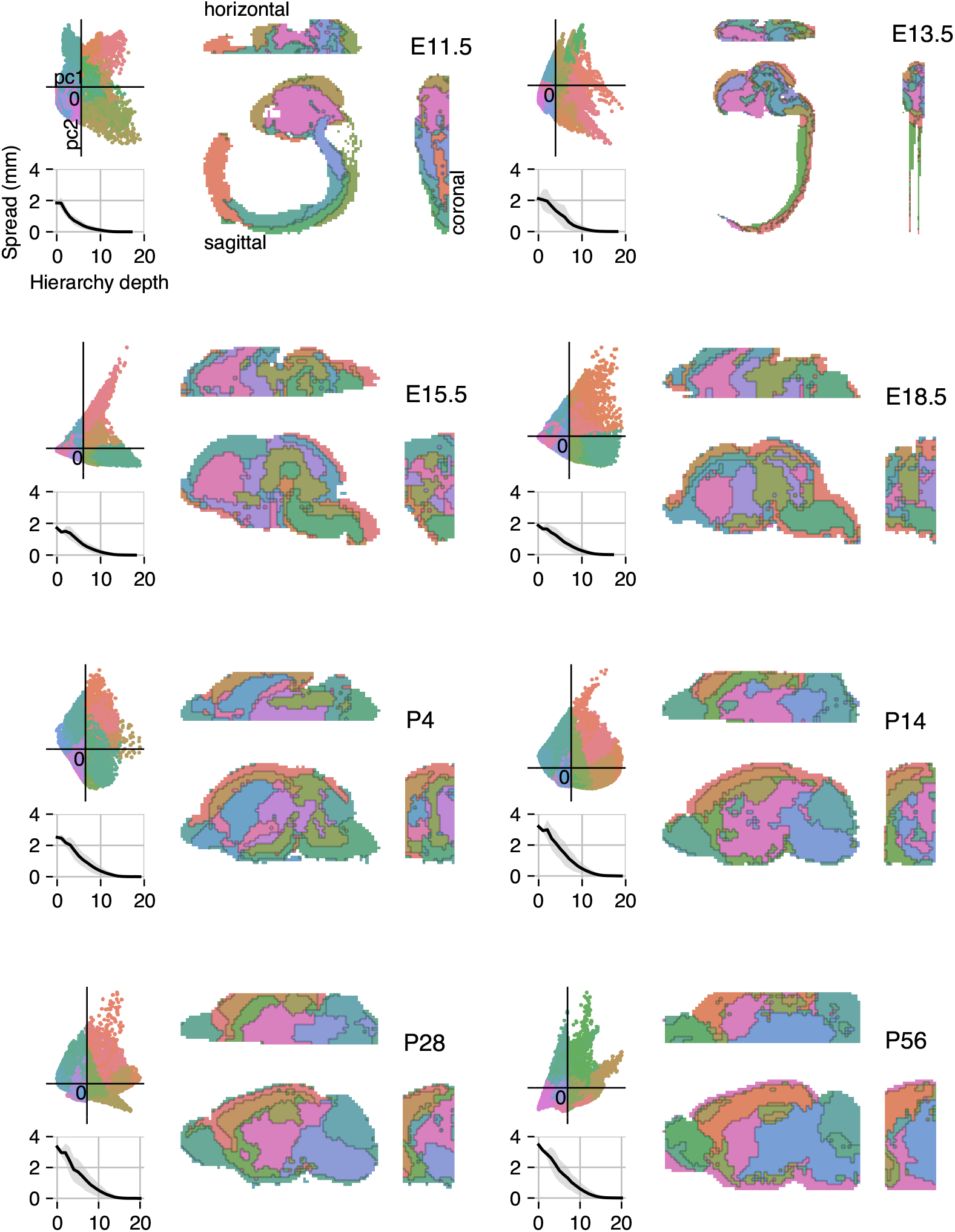
Hierarchical decomposition, as in Figure 5, but for all available time points. Only depth 3—the lowest tile in Figure 5—is shown, but other depths can be inferred by grouping similar colors. Decompositions were performed independently of one another (unlike Figure 6, where established hierarchies are projected across time points). The spatial spread of hierarchical regions goes down with hierarchy depth at each measured time point.

**Figure S12:**
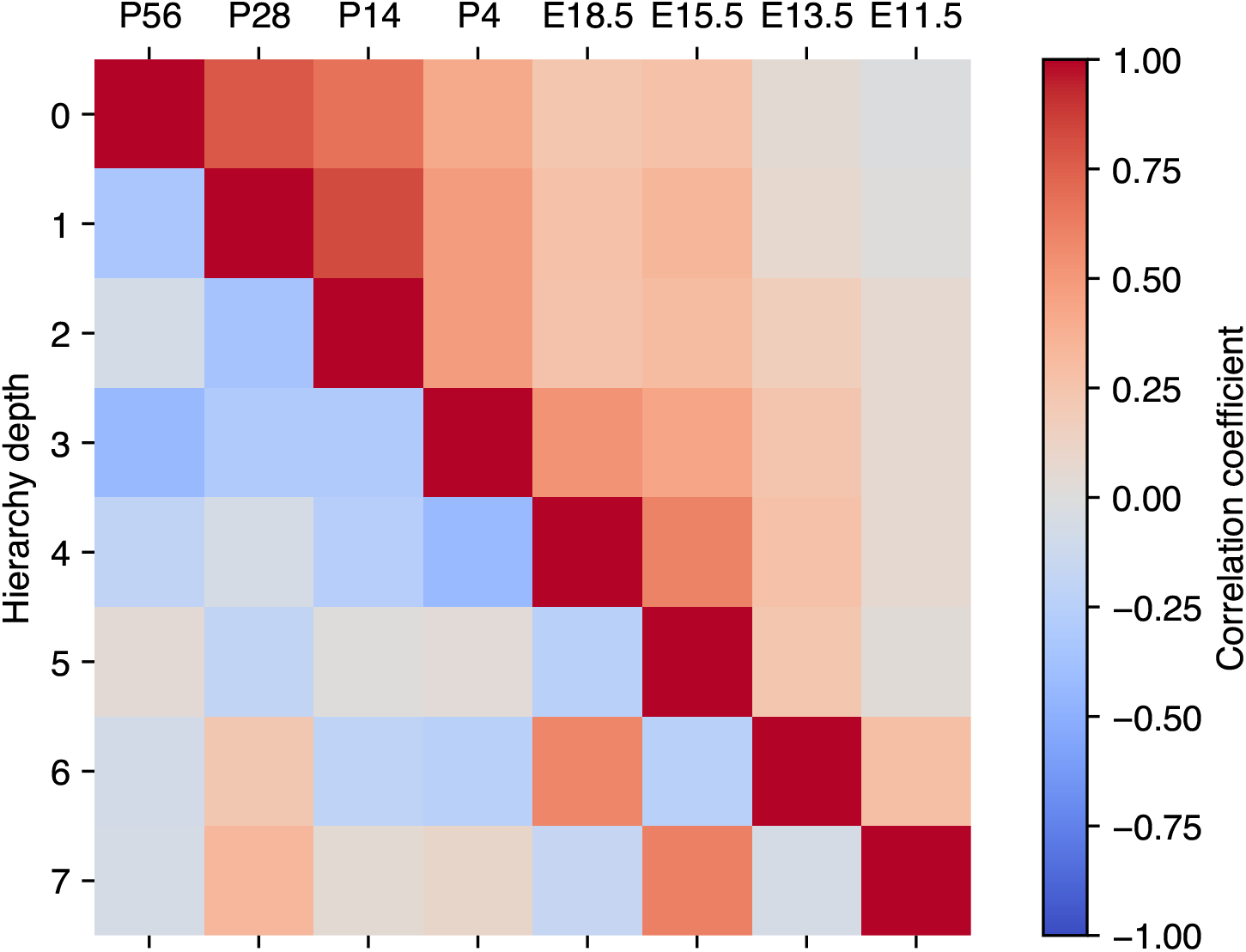
Asymmetry profiles identifying regions of the hierarchical decomposition are poorly correlated within the hierarchy, but correlated across time points. **Upper triangle** Pairwise correlation coefficient between the estimated asymmetries 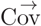 measured at the root of the hierarchies at various time points. Altough the asymmetry measurement is done independently at each time point, the main direction of covariance across all voxels is correlated. Generally, nearby time points are more correlated than distant time points. This correlation is surprising a priori, because the absolute gene expression changes from E11.5 to P56. **Lower triangle** Pairwise correlation coefficient between the estimated asymmetries 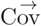 at the root of a hierarchy and other asymmetries within the same hierarchy. (Each column represents a time point, and each row a depth of the hierarchy, with the root at zero depth.) In contrast to standard principal component analysis, orthogonality between components is not enforced by our hierarchical decomposition. Nevertheless, we find that many pairs of components are poorly correlated. This implies that the direction of strongest covariance is not along any single direction for all subsets of voxels, but is rotated in high-dimensional expression space at each iteration of the decomposition. The model assumes that differential gene expression vectors δ, and consequently the asymmetries 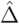 are independent. This matches the observation in the experimental data that the successive 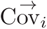 are poorly correlated in expression space. The poor correlation is not by construction, because unlike PCA (Principal Component Analysis), orthogonality is not enforced by our decomposition.

**Figure S13:**
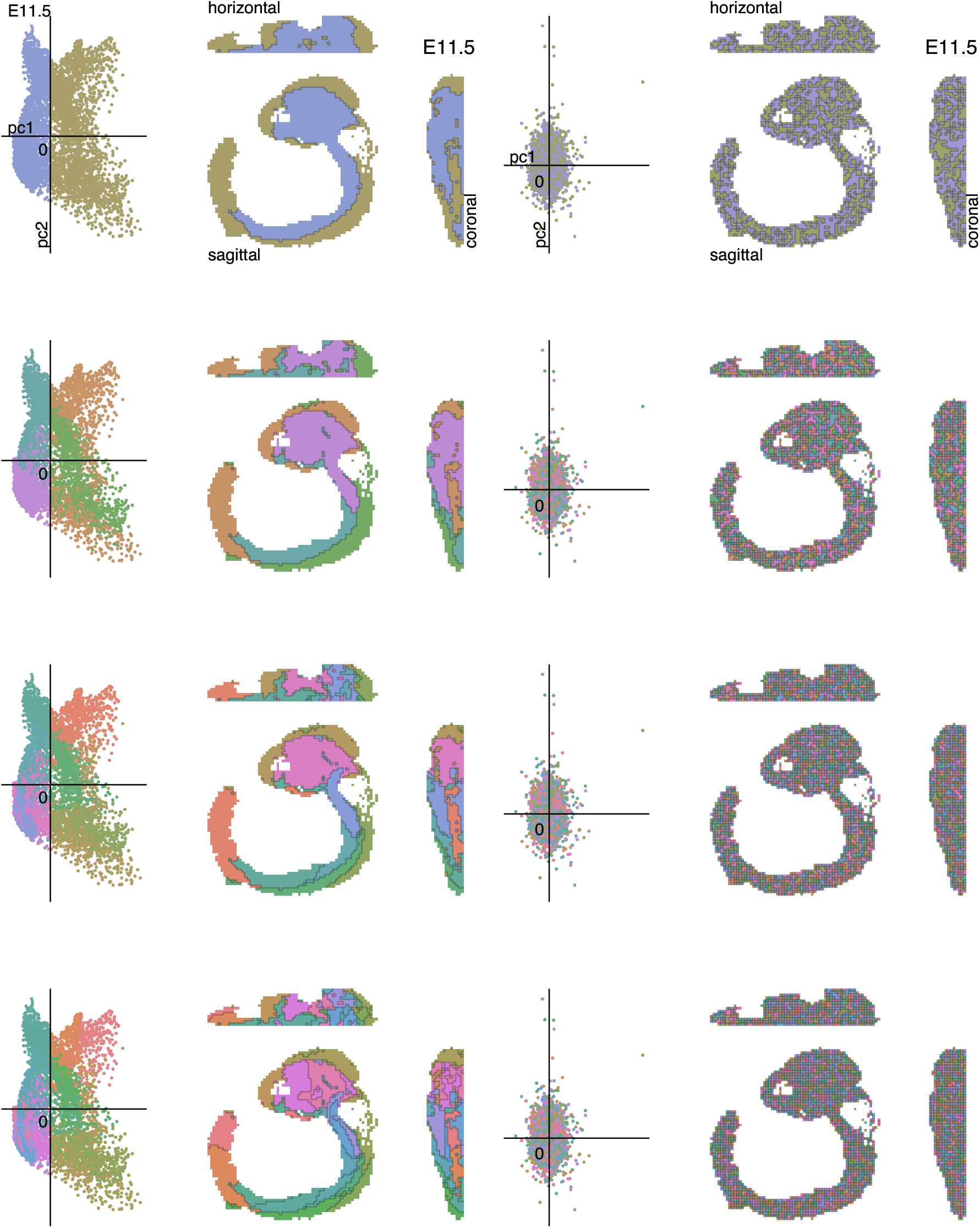
Hierarchical decomposition at E11.5. Analysis and depiction as in Figure 5. Bottom right matrix shows pairwise correlation coefficient among components within the hierarchy at the displayed depths. (Similar to the bottom triangle in Figure S12.)

**Figure S14:**
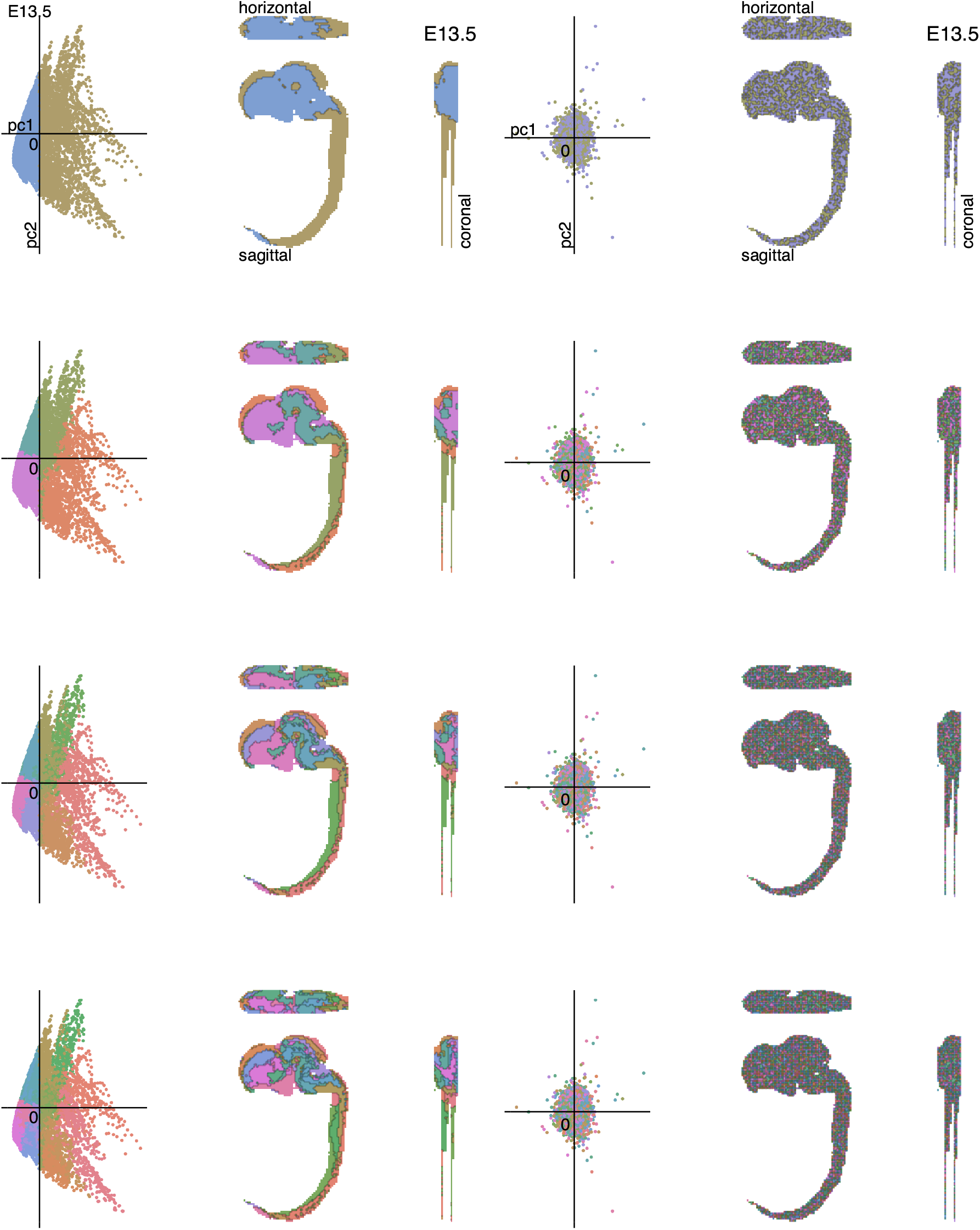
Hierarchical decomposition at E13.5. Analysis and depiction as in Figure 5. Bottom right matrix shows pairwise correlation coefficient among components within the hierarchy at the displayed depths. (Similar to the bottom triangle in Figure S12.)

**Figure S15:**
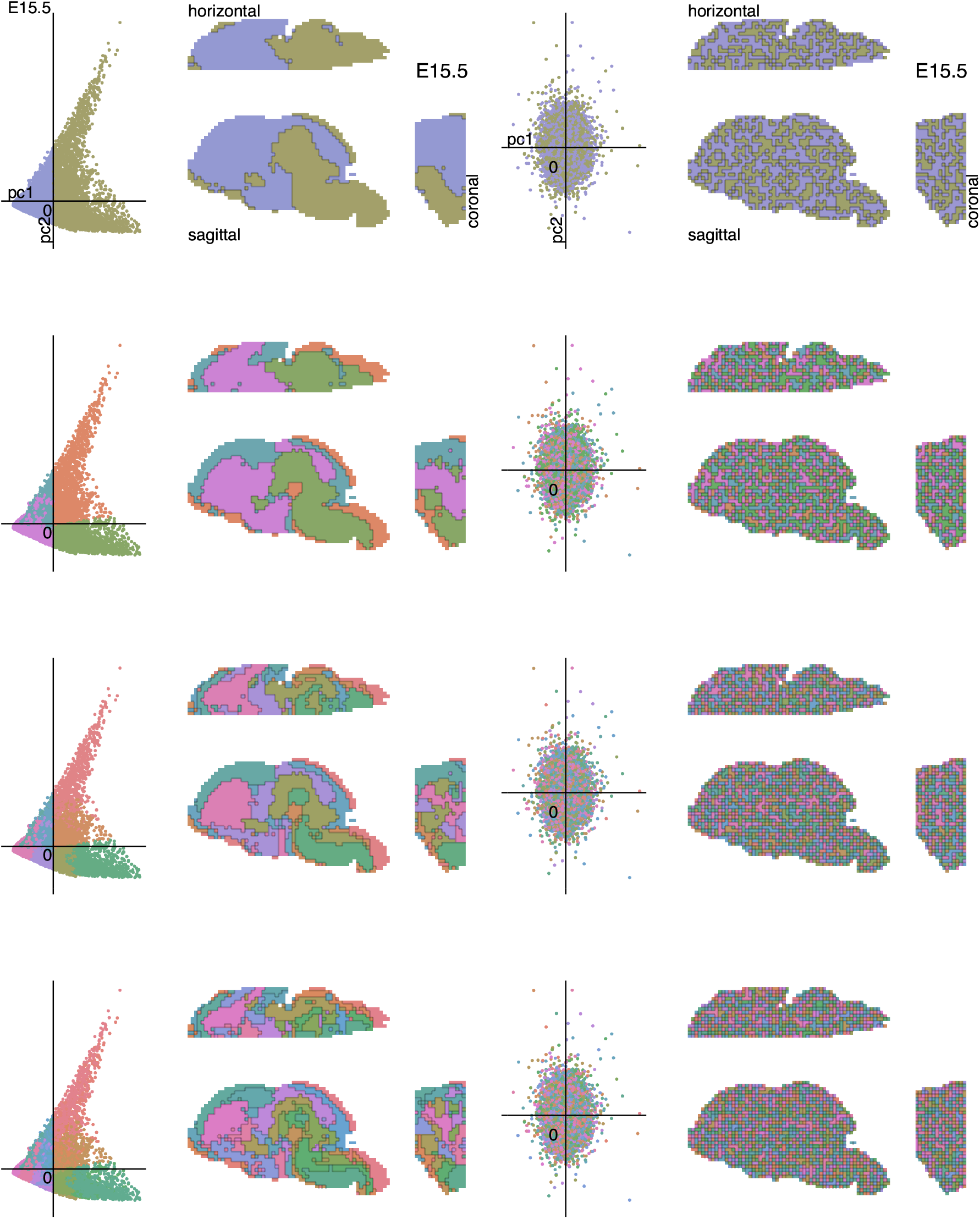
Hierarchical decomposition at E15.5. Analysis and depiction as in Figure 5. Bottom right matrix shows pairwise correlation coefficient among components within the hierarchy at the displayed depths. (Similar to the bottom triangle in Figure S12.)

**Figure S16:**
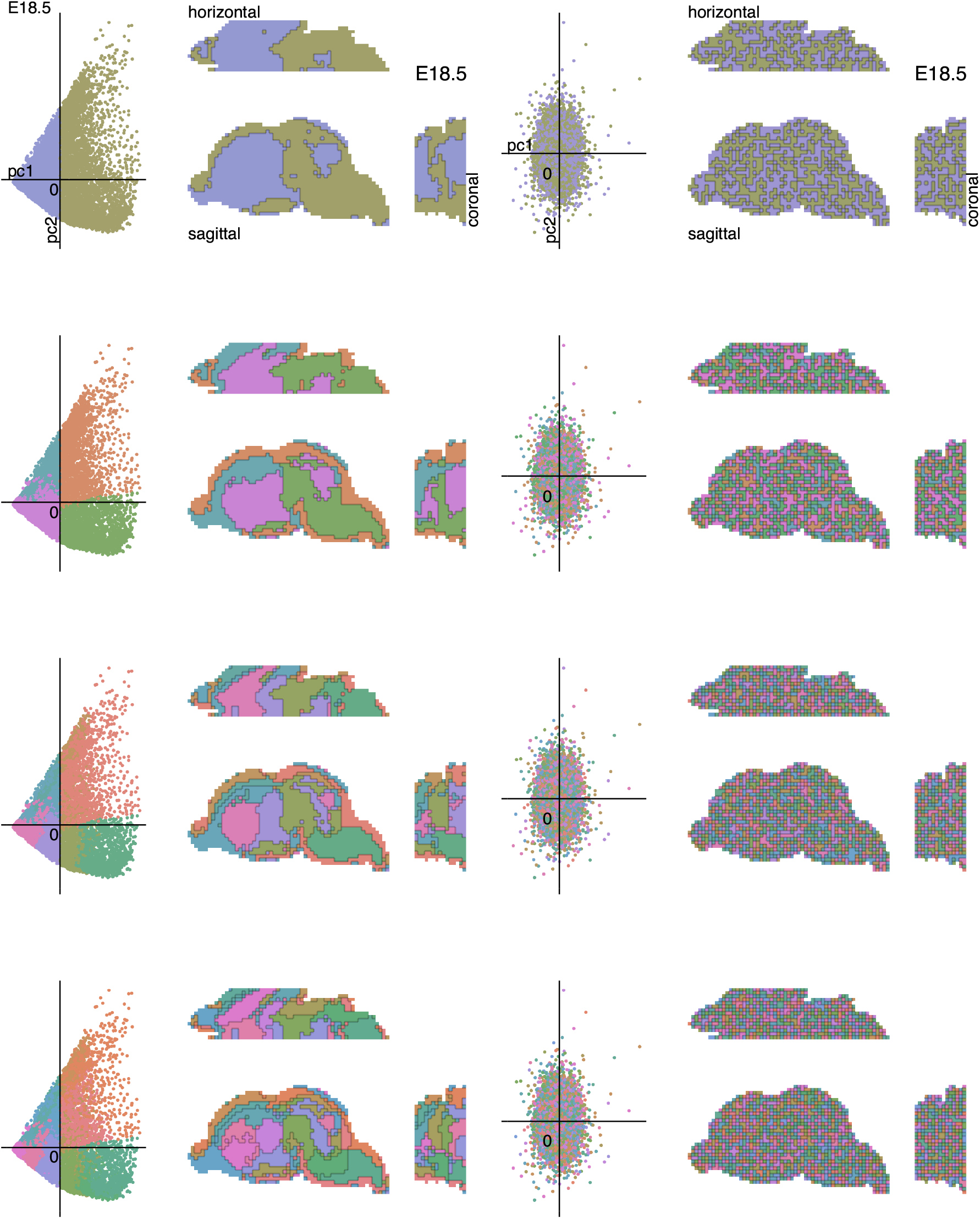
Hierarchical decomposition at E18.5. Analysis and depiction as in Figure 5. Bottom right matrix shows pairwise correlation coefficient among components within the hierarchy at the displayed depths. (Similar to the bottom triangle in Figure S12.)

**Figure S17:**
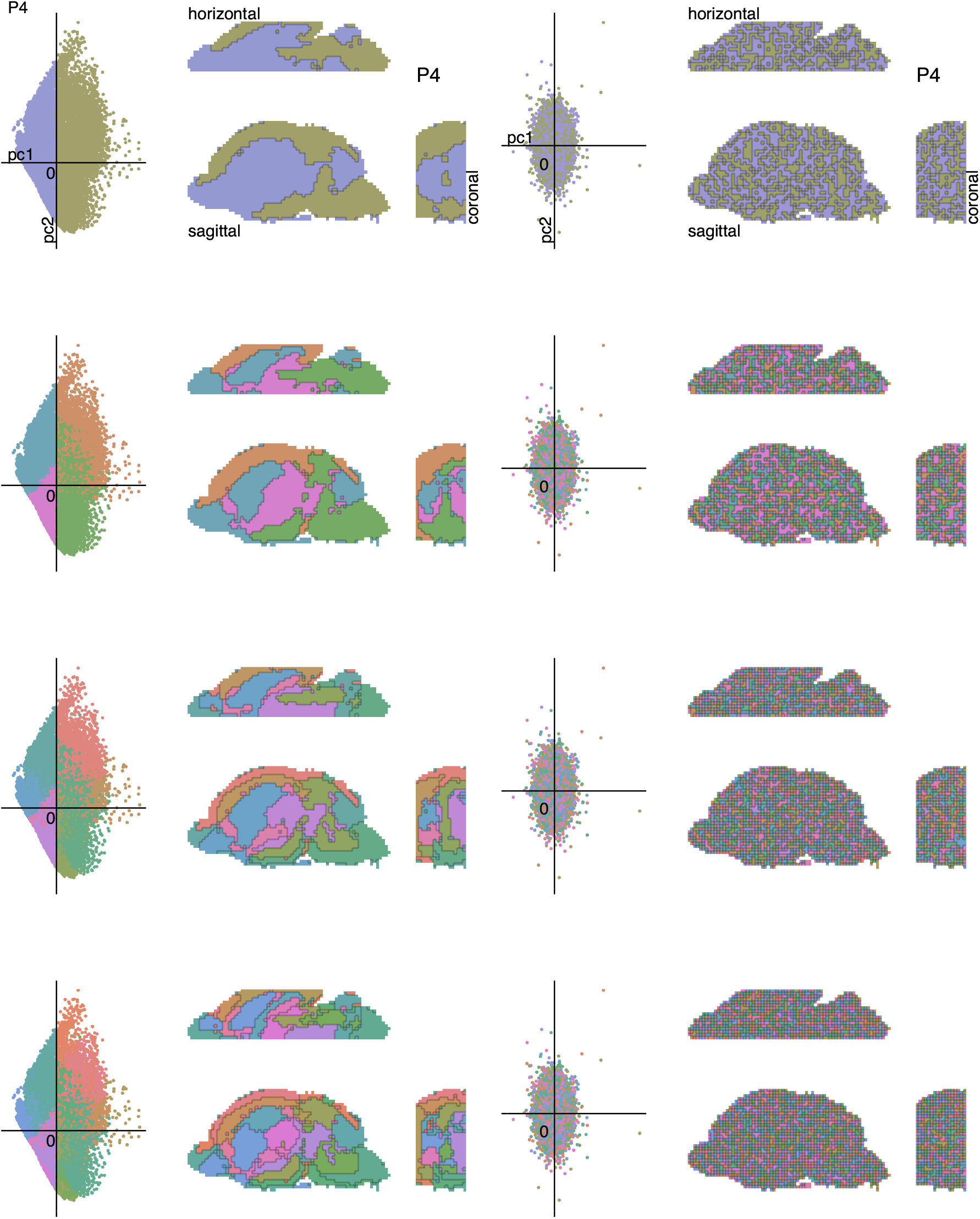
Hierarchical decomposition at P4. Analysis and depiction as in Figure 5. Bottom right matrix shows pairwise correlation coefficient among components within the hierarchy at the displayed depths. (Similar to the bottom triangle in Figure S12.)

**Figure S18:**
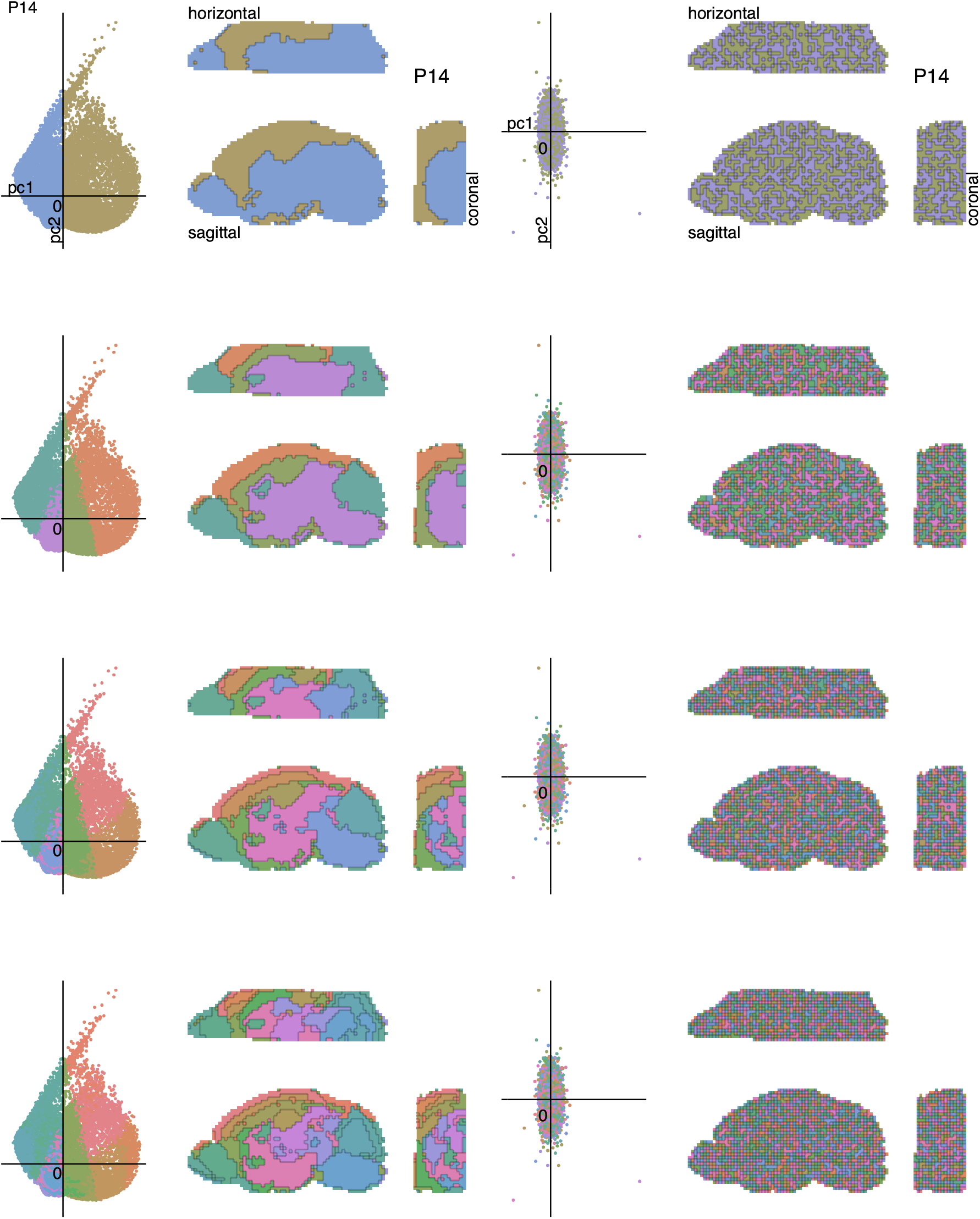
Hierarchical decomposition at P14. Analysis and depiction as in Figure 5. Bottom right matrix shows pairwise correlation coefficient among components within the hierarchy at the displayed depths. (Similar to the bottom triangle in Figure S12.)

**Figure S19:**
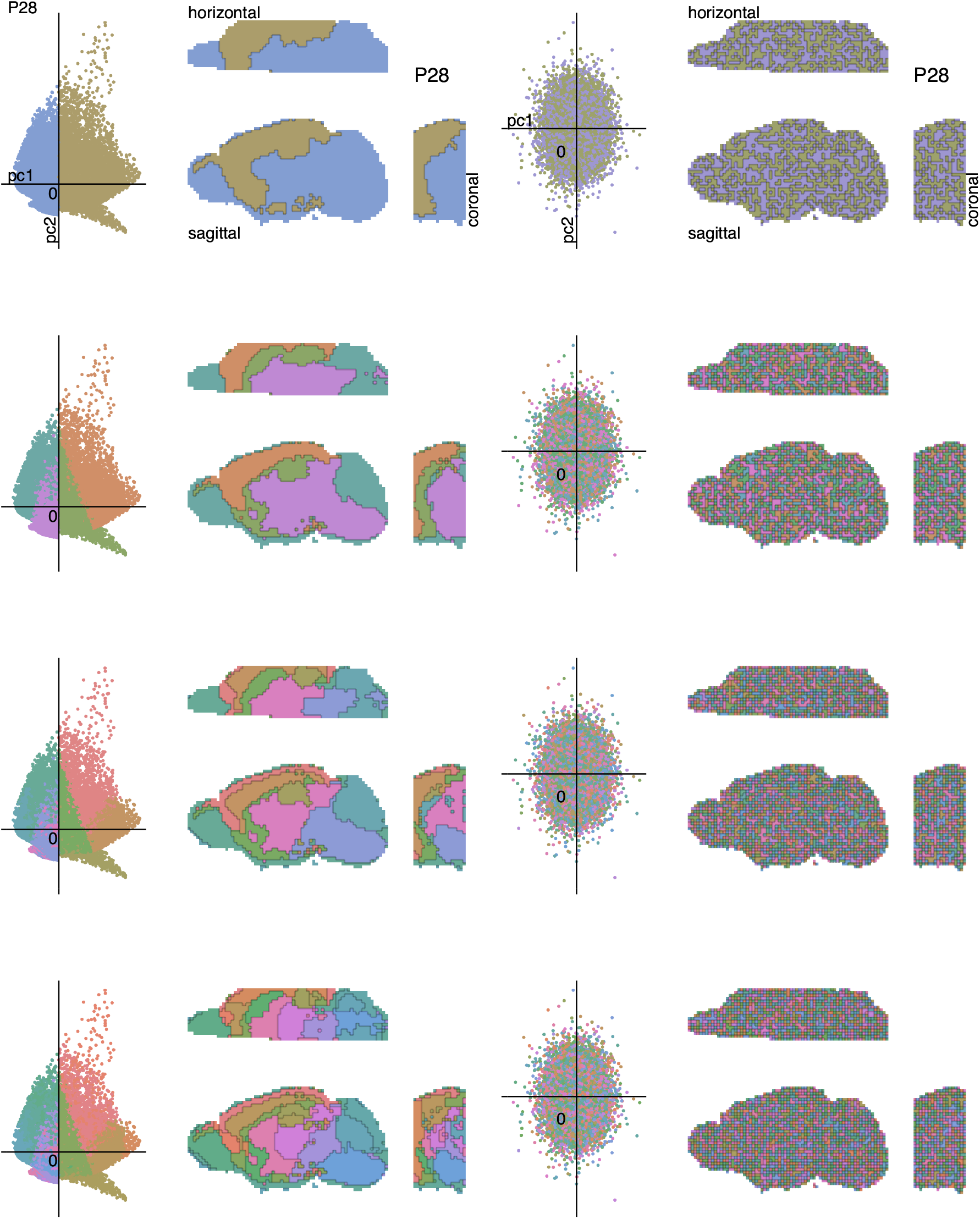
Hierarchical decomposition at P28. Analysis and depiction as in Figure 5. Bottom right matrix shows pairwise correlation coefficient among components within the hierarchy at the displayed depths. (Similar to the bottom triangle in Figure S12.)

**Figure S20:**
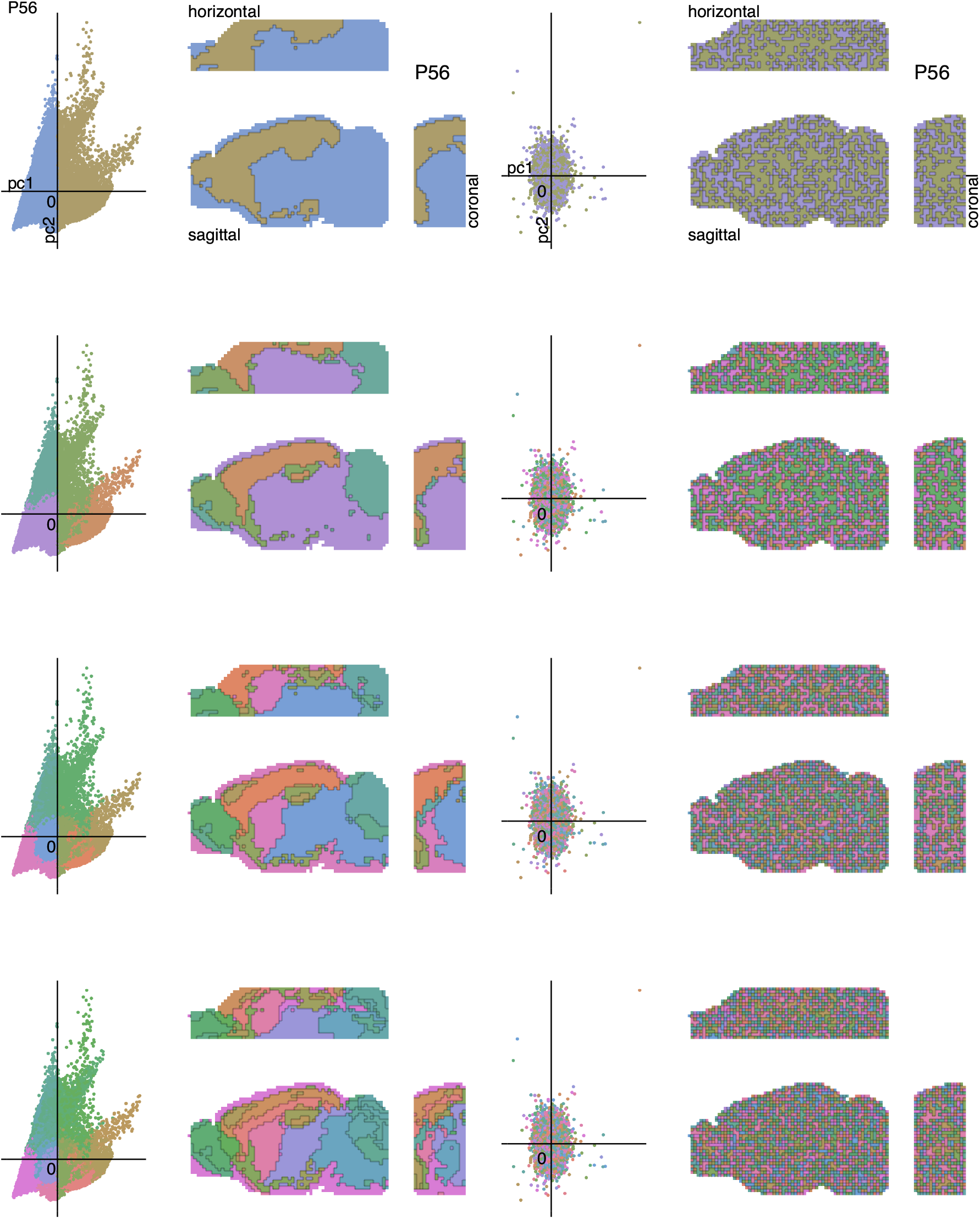
Hierarchical decomposition at P56. Analysis and depiction as in Figure 5. Bottom right matrix shows pairwise correlation coefficient among components within the hierarchy at the displayed depths. (Similar to the bottom triangle in Figure S12.)

## Footnotes

∗ The mouse genome has roughly 2.5 billion base pairs, each encoding 2 bits. At 8 bits per byte this constitutes 625 megabytes, which generously rounds to 1GB. The number for other vertebrates is similar.

